# Regulatory Modules of Human Thermogenic Adipocytes: Functional Genomics of Meta-Analyses Derived Marker-Genes

**DOI:** 10.1101/2021.01.25.428057

**Authors:** Beáta B. Tóth, Zoltán Barta, László Fésüs

## Abstract

Recently, ProFAT and BATLAS scores have been offered to determine thermogenic status of adipocytes using expression pattern of brown and white marker-genes. In this work, we investigated the functional context of these genes. Although the two meta-analyses based marker-gene lists have little overlap, their enriched pathways show strong coincides suggesting they may better characterize adipocytes. We demonstrate that functional genomics of the annotated genes in common pathways enables an extended analysis of thermogenesis regulation, generates testable hypotheses supported by experimental results in human adipocytes with different browning potential and may lead to more global conclusions than single-state studies. Our results imply that different biological processes shape brown and white adipocytes with presumed transitional states. We propose that the thermogenic adipocyte phenotype require both repression of whitening and induction of browning. These simultaneous actions and hitherto unnoticed regulatory modules, such as the exemplified HIF1A that may directly act at *UCP1* promoter, can set new direction in obesity research.

**Highlights:**

- Integrated pathways better characterize brown adipocytes than marker-genes
- Different processes shape the brown and white adipocyte phenotypes
- Thermogenic phenotype may require simultaneous repression of whitening and induction of browning
- Protein network analyses reveals unnoticed regulatory modules of adipocyte phenotype
- HIF1A may regulate thermogenesis by direct control of *UCP1* gene-expression

**Graphical Abstract:** 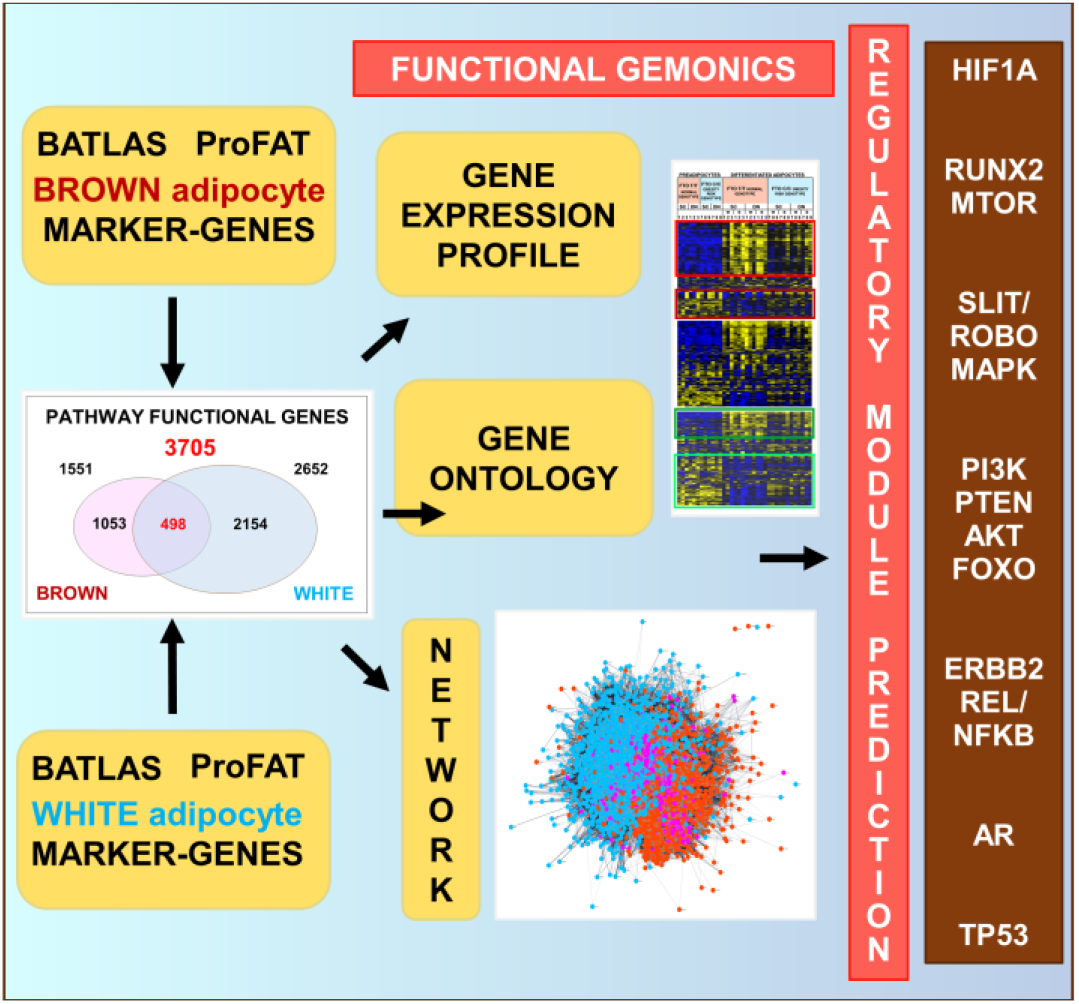

## INTRODUCTION

Directing our bodies to burn excess energy by up-regulating high-energy releasing biochemical reactions at cellular level has become a hot topic for metabolic research, as the obesity pandemic and related pathologies affect much of the population, especially in Western civilizations. The need for a pharmacological treatment is pressing. One promising therapeutic solution to combat obesity is the induction of thermogenesis in adipose tissue (1,2). The number of newly identified genes and proteins that are known to regulate thermogenesis in adipocytes is steadily increasing, yet much is still unknown how to safely and precisely activate adipocyte heat production and clarification of plasticity between thermogenic brown and non-thermogenic white phenotypes is also pending.

The key molecular marker of thermogenicity (browning) in adipose tissue is the expression of Uncoupling Protein 1 (UCP1) which dissipates the proton gradient in the mitochondrial inner membrane (3,4). Further studies have discovered additional important elements (5,6), which, can vary by tissue localisation (7,8,9) and organism type (10,11). In addition, based on genome-wide association studies, a single nucleotide polymorphism (SNP), for example at the locus of the Fat mass and obesity-associated (*FTO*) gene intronic region rs1421085, can critically contribute and modify the development of thermogenic phenotype (12,13). Claussnitzer (2015) demonstrated that the presence of the homozygous C/C obesity-risk alleles in this *FTO* locus results in the expression of nearby genes *IRX3* and *IRX5*, generating less thermogenic adipocyte phenotype. By contrast, in the T/T non-obesity risk (higher thermogenic potential) genotype individuals, the ARID5B repressor is able to bind this DNA site and thus blocks the expression of *IRX3* and *IRX5* allowing the formation of the browning adipocyte phenotype.

Recently, the BATLAS and ProFAT marker-gene scores have been proposed to describe the thermogenic status of a tissue or adipose cell using the expression patterns of specific sets of genes (BATLAS 98 brown and 21 white genes, Suppl. Table 1.(14); ProFAT 53 brown and 6 white genes; Suppl. Table 2.(15)) and hence provide comprehensive, standardised approaches. The two gene-expression based meta-analyses found a distinct molecular signature for the brown and white adipocyte phenotypes and identified basic markergenes for browning prediction. Interestingly, the two sets of genes have little overlap: only 17 common genes can be found (Suppl. Figure 1). However, by examining them using a different approach, we can find more agreement. The majority of both gene sets encode proteins that function in the mitochondria (considering MitoCarta 2.0 database; 16); in fact, 85% and 86% of the browning marker-genes encode mitochondrial proteins in BATLAS and ProFAT, respectively, while none of the white markers do.

In the present study, we intended to investigate the functional context of these two marker-gene sets. Performing Gene Set Enrichment Analyses (GSEA) (17) we found a clear consistency in the enriched pathways of the two brown marker-gene sets, which suggests that these pathways may be critical in generating the thermogenic phenotype. Based on this we further investigated all annotated genes of the enriched pathways, presuming that it may leads to more generally applicable conclusions than single-state studies. To see and analyse the relative expression profile of these genes in adipocytes, we used transcriptional data from our recent study (18) on *ex vivo* differentiated primary human subcutaneous (SC) and deep neck (DN) brown adipocytes, with or without *FTO* rs1421085 intronic SNP (obesity-risk allele). Irrespective to the tissue locality and the applied differentiation protocol (thermogenic induction during differentiation: brown protocol; No thermogenic induction: white protocol; Methods) molecular signature of the *FTO* normal (T/T) genotype adipocytes suggests higher thermognicity then *FTO* obesity-risk genotype (C/C) adipocytes. Therefore, we compared gene-expression pattern based on this approach. Functional genomics of all genes annotated to the identified characteristic pathways of the white or brown adipocyte phenotypes implies that different biological processes shape them, and that downregulation of certain genes could play an important role in thermogenicity. The core module of their interactome points to so far less well-known signaling pathways and transcriptional regulatory elements that may determine adipocyte phenotype and provide appropriate targets in the pharmacological treatment of obesity.

## RESULTS

### GSEA and Network of BATLAS and ProFAT marker-genes reveal biological pathways characteristic for thermogenic adipocytes

First, to see which biological processes and pathways were associated with the basic marker-genes we performed a GSEA separately for the brown and white markers of BATLAS and ProFAT (note that the six ProFAT white genes did not define any enriched pathway; Methodes). Brown marker-genes show high consistency in the enriched pathways: out of the 23 and 23 significantly enriched KEGG (19; Methods) and REACTOME (20; Methods) pathways defined by ProFAT brown marker-genes, 21 and 21 were found also by analysing the BATLAS brown marker-gene set, respectively (Table 1, Suppl. Table 3.A, B, red colour). This suggests that these common enriched pathways are critical in generating the thermogenic phenotype, as the two meta-analyses (BATLAS and ProFAT), which used different databases and selecting algorithms, identified genes from almost the same pathways, albeit largely not the same genes. Importantly, the same enriched pathways defined by the BATLAS and ProFAT brown marker-genes as described above were also found to be significantly enriched based on the preferentially expressed genes in our previous study (18) of the non-obese *FTO* genotype brown adipocytes (Suppl. Table 3 A, B, blue colour), emphasizing their role in shaping the thermogenic phenotype. The pathways defined by the basic brown marker-gene sets were almost completely different from those defined by the BATLAS basic white marker-genes (47 KEGG and 17 REACTOME), as only the Non-alcoholic fatty liver disease (NAFLD) KEGG pathway was identified by both basic marker-genes (Supplementary Table 3 C, D). Furthermore, in both cases the STRING (21) interaction network of the proteins coded by the marker-genes (Figure 1 A,B) show that while there is a high degree of interaction among the brown marker proteins, there is hardly any among the white markers and brown and white markers do not form an integrated network but appear side by side. Taken together, these findings suggest that fundamentally different processes underlie the development of the two adipocyte phenotypes, and that entire pathways may better characterize the phenotype than marker-genes.

**Table 1.** Common KEGG and REACTOME enriched pathways in BATLAS and ProFAT brown Marker-genes. Table show the 21 KEGG and 21 REACTOME pathways that were significantly enriched in both BATLAS and ProFAT brownmarker-gene sets. Pathways listed according to their significances (FDR value) and ordered from highest to lowest.

| Common KEGG enriched pathways in ProFAT and BATLAS Brown Marker-genes | Common REACTOM enriched pathways in BATLAS and ProFAT Brown Marker-genes |
| --- | --- |
| Metabolic pathways | The citric acid (TCA) cycle and respiratory electron transport |
| Citrate cycle (TCA cycle) | Metabolism |
| Carbon metabolism | Pyruvate metabolism and Citric Acid (TCA) cycle |
| Thermogenesis | Respiratory electron transport, ATP synthesis by chemiosmotic coupling, and heat production by uncoupling proteins. |
| Parkinson's disease | Citric acid cycle (TCA cycle) |
| Huntington's disease | Respiratory electron transport |
| Oxidative phosphorylation | Complex I biogenesis |
| Non-alcoholic fatty liver disease (NAFLD) | Mitochondrial Fatty Acid Beta-Oxidation |
| Alzheimer's disease | Fatty acid metabolism |
| Biosynthesis of amino acids | Glyoxylate metabolism and glycine degradation |
| 2-Oxocarboxylic acid metabolism | mitochondrial fatty acid beta-oxidation of saturated fatty acids |
| Fatty acid metabolism | Regulation of pyruvate dehydrogenase (PDH) complex |
| Retrograde endocannabinoid signaling | Beta oxidation of hexanoyl-CoA to butanoyl-CoA |
| Pyruvate metabolism | Beta oxidation of decanoyl-CoA to octanoyl-CoA-CoA |
| Fatty acid degradation | Signaling by Retinoic Acid |
| Valine, leucine and isoleucine degradation | Metabolism of lipids |
| Fatty acid elongation | Mitochondrial protein import |
| Glyoxylate and dicarboxylate metabolism | Lysine catabolism |
| Glycolysis / Gluconeogenesis | Metabolism of amino acids and derivatives |
| Cardiac muscle contraction | Branched-chain amino acid catabolism |
| PPAR signaling pathway | Transcriptional activation of mitochondrial biogenesis |

**Figure 1.**
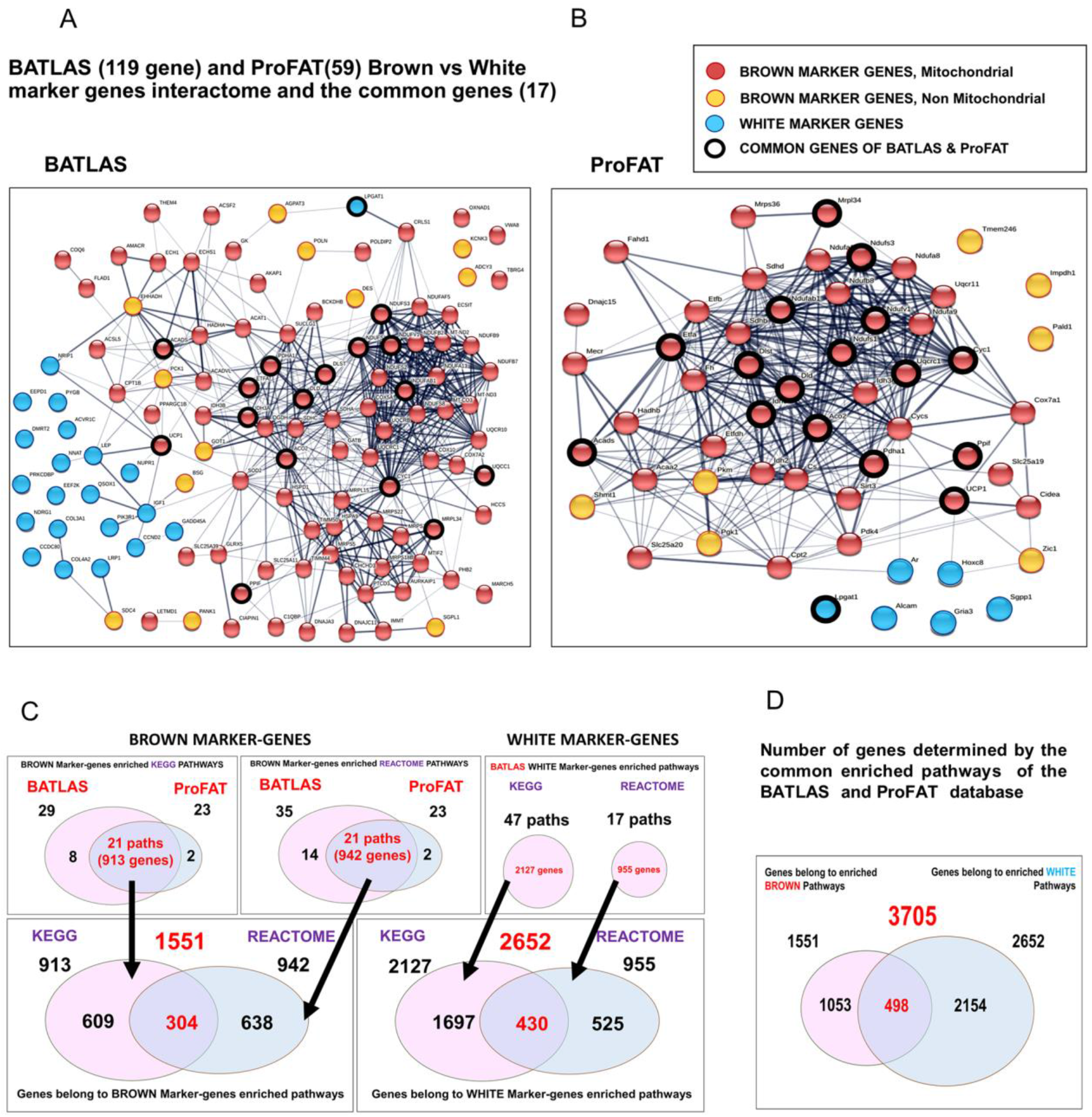
Interactome, Relative gene-expression and Pathway analyses of the BATLAS and the ProFAT marker-genes. Interactome map of **(A)** BATLAS and **(B)** ProFAT brown and white marker-genes; interaction score confidence value: 0.4 **(C)** Venn diagrams show the pipeline of the expanded gene-set generation; the number of the KEGG and REACTOME pathways enriched by the marker-genes from BATLAS and ProFAT database and the number of the white and brown pathway associated genes. **(D)** Venn diagram summarizes the number of genes determined by the enriched pathways of the BATLAS and ProFAT database.

### Considering all genes from the enriched pathways of the BATLAS and ProFAT marker-genes provides an expanded gene set to explore regulation of adipocyte thermogenicity

Considering the high consensus of the identified pathways of BATLAS, ProFAT and the preferentially expressed genes of the FTO non-obese genotype brown adipocytes, we can reasonably assume that all genes belonging to the commonly enriched brown and white pathways may play a role in the formation of the adipocyte phenotype depending on the particular condition. This prompted us to collect all genes associated with the BATLAS and ProFAT brown and white pathways and continue the systematic analysis with these expanded gene sets to better explore potential regulatory processes in thermogenesis and understand the special relationship between genes forming the brown (thermogenic) and the white (non-thermogenic) adipocyte phenotype. The implemented pipeline for generating the expanded brown and white gene-set is summarised in Figure 1C and D (see also in the methodology section). Altogether 3705 genes, 1551 brown, annotated to 44 pathways (brown pathway genes) and 2652 white, annotated to 64 pathways (white pathway genes) were identified by the enriched pathways derived from BATLAS and ProFAT marker-genes when KEGG and REACTOME analyses results were combined (Figure 1C, D; Suppl. Table 4-7). The KEGG and REACTOME pathway analyser use different approaches, as only a minority of the genes were identified by both (304 in the case of the brown and 430 in the case of the white marker-genes). Although the basic brown marker-gene set defined pathways were different from those defined by the basic white marker-genes, there were overlaps between the annotated genes of the pathways (Figure 1D). These 498 common genes may confer an interface between the two phenotypes balancing the regulation of the thermogenic capacity of the adipocytes, hence we named them linker genes/proteins. Not all of the 3705 identified genes (expanded gene set) are expressed in adipocytes based on our study with neck adipocyte, but they can be involved in the higher-level regulatory processes of thermogenesis, therefore we worked with all of these genes in the following network analyses; however, the following heatmaps of the gene-expression profile only depict genes that were expressed in adipocytes from our previous study (18) to increase the clarity of our visualisations.

### The predictive role in thermogenicity of the expanded gene sets is supported by the expression pattern in differentiated adipocytes

To see how genes in the expanded sets are involved in the formation of a particular adipocyte phenotype, their relative expression profiles are shown on three separate heatmaps, namely Brown (1053), Linker (498) and White (2154) genes, as they are manifested according to the *FTO* obesity-risk genotypes in the above-mentioned study with human neck adipocytes (18) (Figure 2A). The majority of the genes belonging to the Brown pathways have higher expression in the more thermogenic samples (red box): 540 genes, e.g. *DIO2, FABP3-4, PDK2-4, PC, CKMT1A, ELOVL3, DHRS9, LPL* and *PLIN1,2,4,5*. On the other hand, most of the White genes are more highly expressed in the less-thermogenic samples (green box): 1211 genes, e.g. *IGF2, HAPLN1, NOX4, COMP, IL11, VCAM1, ID1, LEFTY2* and *ADRA2C*. These genes may be directly and positively involved in the development of the given phenotype. A smaller portion has an opposite expression profile, suggesting a negative regulatory role in the pathway shaping a given brown (dark-red box) or white (dark-green box) phenotype. There are also genes, that show no differential expression profile. This could be because the activity of the proteins encoded by these genes are not regulated at the gene expression level, but rather via post-translational modification or splicing variation which can allow a much quicker response to changing conditions. It is also possible that a smaller change in expression of these genes results in a greater effect due to their position in the signaling network. The Linker genes show higher (e.g. *HK2, PGC1A and B, LIPE, SOD2, CD36, LDHB, PDK1* and *CPT1B*) or lower (e.g. *GNAO1, GNG7, ADCY4, BDNF, SCD5, GNB3, CPT1A* and *TGFB1*) expression in equal proportions in samples of *FTO* normal genotypes, which may indicate an important regulatory role in both directions to switch on/off thermogenesis. Taken together, the expression profile of the expanded gene set largely follows that of the basic marker-genes (Suppl. Figure 2 A. B). However, since it encompasses complete biological processes, unlike the marker-gene sets, which typically only highlights positive regulatory participants, it also identifies those involved in negative regulation providing a suitable basis for a more complex analysis of adipocyte thermogenicity.

**Figure 2.**
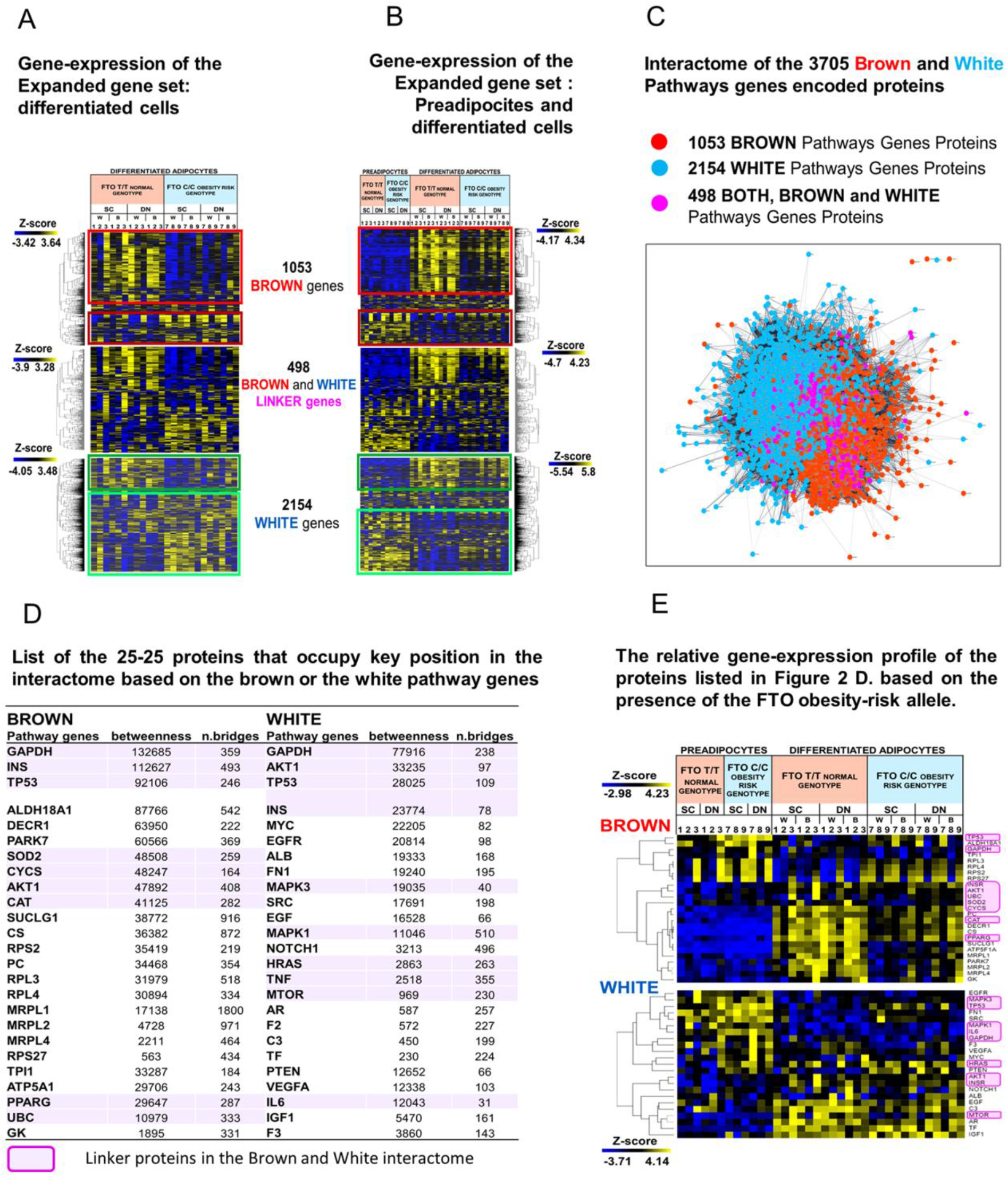
Gene-expression profile and interactome analyses of the Expanded gene set from the Brown and White pathway genes. **(A)** Heatmaps show separately the relative gene-expression profile of the genes enriched in the Brown, the White and both the Brown and White pathways (Linker genes) in differentiated adipocytes and **(B)** in preadipocytes and differentiated adipocytes based on the presence of FTO obesity-risk alleles; we used samples with *FTO* T/T-risk-free (donor 1-3) and C/C-obesity-risk alleles (donor 7-9) and left out the *FTO* T/C heterozygous samples (donor 4-6) which are shown for some cases in the paper Tóth et al., 2020. B:Brown differentiation; W: White differentiation; SC: Subcutameous; DN: Deep-neck **(C)** Interactome of the 3705 Brown and White Pathways genes encoded proteins; Interaction score confidence value: 0.4. **(D)** List of 25-25 proteins that occupy a pivotal position in the interactome based on the betweenness centrality score (betweenness) and the number of bridges (n.bridges) from network analyses of the Brown or the White pathway genes. Magenta Highlights the Linker-proteins that occupy a pivotal position in the interactome based on the Brown or White pathway genes coded proteins. **(E)** The relative gene-expression profile of the 25-25 proteins with highest betweenness centrality score and number of bridges from the Brown and White pathway genes based on the presence of the FTO obesity-risk allele. Magenta highlights the linker proteins.

### Comparison of pre- and differentiated adipocyte transcriptomes uncovers the importance of gene-suppression that may regulate thermogenesis

Gene expression patterns of adipocyte progenitor cells are usually not reported in studies addressing thermogenic potential, but it may be important to examine them as well, to see not only the differences between the matured adipocytes but the direction of the regulatory processes too. Most of the genes of the expanded gene set show differential expression after differentiation. When preadipocytes are also considered, the expression difference between the *FTO* normal and obesity-risk samples was less pronounced due to the large difference between preadipocyes and differentiated cells (Figure 2A, B, red and green boxes; note: similar expression profiles are presented by the BATLAS and ProFAT basic marker-genes themselves when preadipocytes were also included in the analysis, Supplementary Figure 2 C.D). The main expression pattern of the differentiated cells remained the same. However, comparison to preadipocyte gene-expression reveals that many of the Brown genes expressed higher in the *FTO* normal brown adipocytes after differentiation are also expressed in obesity-risk samples, albeit to a lesser extent (Figure 2B. red box). This indicates that the two genotype samples only differ in the induction level of these genes and they are not actually suppressed in obesity-risk genotype samples. This emphasise the importance of proper controls. On the other hand, considering the cluster of Brown genes downregulated during differentiation (Figure 2 A, B dark red box) these genes are indeed highly repressed in the normal *FTO* genotype brown adipocytes, especially when compared to preadipocytes, and the difference between the two *FTO* genotypes reflects the extent of repression. This highlights to the possible importance of repressed genes which so far have received less attention in analyses of the thermogenic processes. The revealed 175 genes repressed during differentiation include *LTC4S, HACD4, LHPP, PRDM16, GPX3, GPX8, NNMT* (Suppl. Table 8). The White genes, which are mostly suppressed in *FTO* normal brown adipocyte samples in the neck area after differentiation, are needed to form the white phenotype and their presence could interfere with thermogenesis (Figure 2B. green box). Upregulated white genes in *FTO* normal genotype brown adipocytes could be negative regulators of the white phenotype and not directly involved in the thermogenic processes (eg.: *TGFA, LPAR3, SLC2A4, MGST2, FGF1, LPIN1, DUSP6* and *CSF2RA*); however, increases in their expression level may repress the white phenotype, opening the way for appropriate browning. As a conclusion, comparative analysis of differentiated adipocyte samples based on the relative gene expression profile can explore regulatory processes in more detail when the status of preadipocytes is also included. It suggests that in addition to the upregulated genes, the downregulation of certain genes or pathways could have a critical role in determining adipocyte cell fate.

### Analyses of the interactome of the encoded proteins of the brown and white expanded gene sets identifies key participants in network integrity and new regulatory elements in adipocyte thermogenesis

To determine which functional modules and regulatory elements play a central role in the complex process of adipocyte phenotype formation, we mapped the interaction network of proteins encoded by the expanded gene set, as all of these proteins are potential participants. The number of interactions is very large between the products of either the brown or the white genes of the expanded gene sets (Figure 2C), which was expected, since genes of specific pathways make up the expanded brown and white gene sets and interactions level are usually high between genes that belongs to one particular pathway. For that reason, the network shows a so-called “hairball” profile (22), in which the sub-clusters are not visible, but the brown (red) and white (blue) genes/proteins are spectacularly separated, as was the case of the basic ProFAT and BATLAS marker-gene set (Figure 2C; Figure 1D, E; note: similar network appeared, when interaction confidence score was 0.7, not shown). Unsupervised Markov CLustering Algorithm (MCL) was used to explore the sub-clusters (Cytoscape) and the majority (41 (73%) out of 56 clusters which contain more than 8 proteins) of them contains either BROWN and LINKER or WHITE and LINKER proteins, while only 27% have exclusive BROWN and WHITE proteins mixed (Suppl. Figure 3; Method). These further strengthens our conclusion that mainly different processes shape the development of the brown and white adipocyte phenotypes and we can presume possible transitional states. Therefore, inhibition of the formation of one phenotype would not necessarily imply induction of the other one and *vice versa*. Based on this, we propose that the healthy induction of the thermogenic adipocyte phenotype may require simultaneous repression of whitening and activation of browning.

It can be also observed that genes/proteins common to both brown and white pathways are more likely to occur in the middle of the network landscape (Figure 2C, magenta, 498 linker genes/proteins) crossing the interface of the brown and white parts rather than coinciding with it. This may mean that some of these linker proteins, not only connect the two phenotypic processes, but are also deeply embedded in the networks of one or other phenotype, and thus have the potential to have a deciding effect on determining the particular phenotype. To identify them, a separate interactome analysis of the brown and white pathway genes/proteins was performed when the linker proteins were involved in both: 25-25 proteins with the highest Betweenness Centrality score and Bridges Number are shown (Figure 2D) highlighting linker proteins (magenta). These proteins possess the potential of contributing to form either the brown or white phenotype, but of these, linker proteins may be the ones that could connect and balance these two functions (magenta). Their expression profile (Figure 2E; upper heatmap) suggests that among the brown pathway genes only *TP53, ALDH18A1, RPL3-4, RPS2* and *27* are repressed in the *FTO* normal brown adipocytes, pointing to the importance of their downregulation in brown phenotype formation, while the majority of these genes show higher expression in the *FTO* normal samples indicating a possible positive regulatory role or that is the consequences of browning (eg.: *PPARG, SUCLG1, PARK7*). On the other hand, from the white genes *SRC, MAPK3* and *FN1* show increased expression in the obesity-risk brown adipocytes (Figure 2E; Lower Heatmap) and these genes may decrease the themogenic processes. Interestingly, four from the linker proteins (GAPDH, INS/INSR, TP53 and AKT1) appear as key in maintaining the interactome integrity in both the expanded brown and white gene networks, as they appeared in both lists, emphasising their pleiotropic function in biological processes forming the thermogenic phenotype.

In another approach, the expanded gene set derived from brown and white pathways (3705 genes) was considered as a whole, with the assumption that all of them could have a role (either positive or negative regulation) in determining of the adipocyte phenotype. We have identified 30 proteins that bridge the largest number of functional units (clusters) in the network (Figure 3A). These have potentially the greatest impact on others in the network, therefore we considered them as the central pillars (core-module proteins) and further investigated their network relations, expression profile and function. Most of them belong to both the brown and white pathways (linker genes/proteins; Figure 3A. magenta) and are also closely related to each other, as shown by their interactome (Figure 3B). In addition, the linker proteins are aggregated in the middle, further supporting their possible central role in the formation of adipocyte phenotypes, as they can connect the distinct sub-processes that could take place in the differentiated adipocytes. According to KEGG and REACTOME pathway analysis, these 30 genes belong to a large number of pathways (the five most relevant ones are shown on the right side of the interactome figure (Figure 3B)), which further highlights their versatile role. The HIF1A and PIP3-AKT signalling pathways and PTEN regulation are the most significant modules, while the mTOR, MAPK, SLITs-ROBOs, ERBB2, and RUNX2 signalling pathways are also highly enriched by these genes/proteins. The functions that can be related to them were explored by GSEA and the regulation of gene expression, the protein metabolic processes, the catalytic activity and the cell death emerged as the most significantly enriched biological pathways/processes. The Core-module protein gene expression profiles show that some of them are highly expressed in *FTO* normal samples and may be positive effectors, while others show lower expression and can be involved as negative regulators in the process of differentiation to thermogenic adipocytes (Figure 3C.). Thus, interaction analysis of the expanded gene set emphasizes that different biological processes shape the two adipocyte phenotypes and that linker genes could play a central role in this, especially some pleiotropic genes that can regulate both brown and white differentiation processes according to the particular condition.

**Figure 3.**
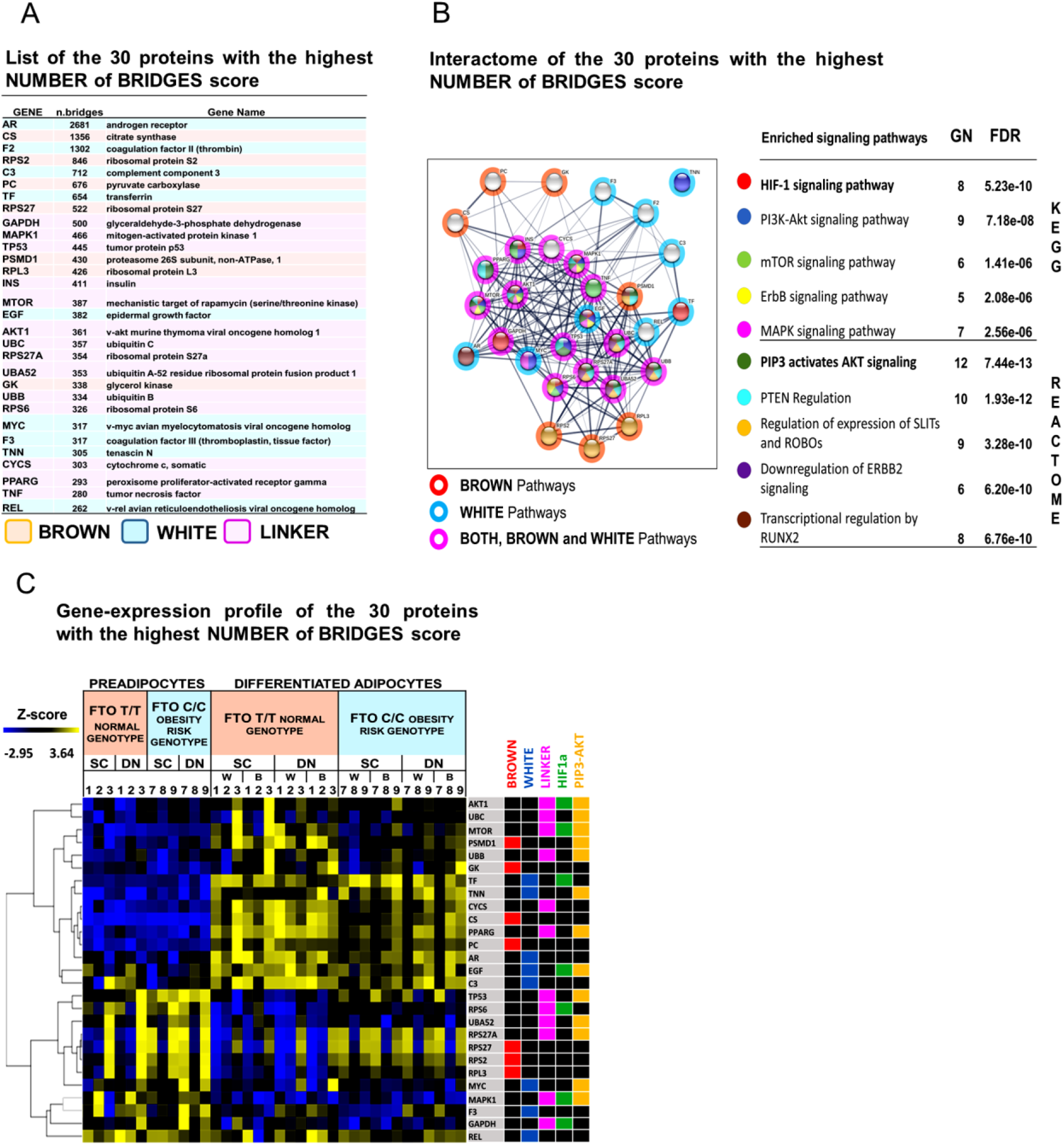
List, Interactome analysis and relative gene-expression profile of the 30 proteins with highest number of bridges in the network of the Expanded gene set. **(A)** List of the 30 proteins with the highest NUMBER of BRIDGES score (core module-proteins) based on the interactome network analyses of the expanded gene set coded proteins (3705 genes) **(B)** Interactome of the 30 core-module proteins, colour of nodes marks the enriched pathways and the colour of rings shows the type of the gene set the gene belongs; red: brown gene-set, blue: white gene-set, magenta: linker genes; right panel lists the enriched signaling pathways, the number of genes (GN) belonging to this pathway from the 30 core-module proteins and the significance level of the enrichment with False Discovery Rate value (FDR). **(C)** Gene-expression profile of the 30 core-module proteins, right panel marks genes that belong to expanded Brown gene set (red), expanded White gene set (blue), Linker genes (magenta), HIF1A pathway (green) and PIP3-AKT (yellow) pathway.

### Interactome of the expanded gene set and the analysis of the UCP1 promoter reveal potential transcriptional regulatory elements such as HIF in adipocyte thermogenesis

Gene expression regulation was one of the significantly enriched pathways by the 30 core-module proteins, so we examined which Transcription Factors (TFs) appeared among them and *in silico* investigated whether they could directly regulate the gene expression of UCP1, the major thermogenic protein. MYC, REL, PPARG, TP53 and AR are among the 30 core-module proteins, so they can directly affect the expression of many other genes and thus may have a broad-spectrum effect (Figure 3A). Furthermore, according to the Eukaryotic Promoter Database (EPD) (https://epd.epfl.ch) (23), they have their binding site sequence (Response Element; RE) in the *UCP1* promoter and/or enhancer region (−5MB-TSS; p <0.0001; TP53 p <0.001, Figure 4A). We also investigated the three transcription factors that emerged in the significantly enriched signalling pathways, namely HIF1A, FoxO1 and RUNX. Intriguingly, they also have their binding sequence in the upstream DNA element of the *UCP1* and the enrichment of the HIF1A RE has the highest probability (p <0.00001; others p<0.0001) (Figure 4A). The role of HIF1A in obesity has been emphasized in hypoxic conditions but less studied in normoxic environments (24). Although HIF1A expression is already reported in thermogenic adipocytes even in UCP1 KO mice (25, 26). That suggests, that the presents of HIF1a could not be the only consequence of termogenic activation induced UCP1 expression connected physiological changes, such as increased ROS production (26). Furthermore, it is not fully clear whether a hypoxic condition is present during thermogenic induction or a pseudo-hypoxic condition results the stabilised HIF1A, and what function is associated with the stabilised HIF1A. To investigate the functionality may connected to the HIF1A presence in thermogenic adipocytes we checked published chip-seq. data, however, we found no HIF1A or HIF1B (ARNT) chip-seq. experiment in thermogenic adipocytes in neither the ChIP-Atlas (http://dbarchive.biosciencedbc.jp; 27) nor in the ChIPSummitDB database (http://summit.med.unideb.hu;28). For other cell types or conditions, *UCP1* was not among the target genes of HIF1A or HIF1B (ChIP-Atlas), or there was no experimental evidence for HIF1A-HIF1B binding to the *UCP1* promoter (TSS-5KB) (ChIPSummitDB). However, this HIF1A binding motif region is an active regulatory element of the *UCP1* promoter, as other TFs bind to this DNA part (Figure 4B; ChIPSummitDB), and the histone methylation and acetylation state also suggests that (Suppl. Figure 4A; https://genome-euro.ucsc.edu). The nucleotide sequence of the HIF1A-HIF1B RE and its immediate vicinity in human is also found in the homologous promoter region of the paralog *UCP2* gene (Figure 4C; EPD database; Suppl. Figure 4B) which is evolutionarily the closest descendant to *UCP1* (29). In addition, both the ChIP-Atlas and the ChIPSummitDB database show, based on experimental data, that the HIF1A-HIF1B heterodimer does bind to this part of the *UCP2* gene promoter (Figure 4D). According to the comparative genomic analysis of the UCSC genome browser, this is a phylogenetically new response element appearing only in primates and is not found in the *UCP1* or *UCP2* promoter of non-primates mammals such as mouse or dog (Suppl. Figure 4 A,B,C) which makes *in vivo* study difficult. Addition to this, Figure 4E. shows the schematic representation of the *UCP1* promoter region (−250--285) with the identified putative binding site of HIF1A-HIF1B (−263) and many overlapping and proximal binding sites for other transcription factors enriched in this region as revealed using the EPD database (p<0.0001). The emerging picture shows an extensive group of potential transcription factors that may alternate or co-interact to regulate transcriptional activation or repression of *UCP1* at this region and can directly modulate the thermogenic activity of the adipocytes according to actual needs and circumstances. HIF1A is a basic helix-loop-helix motif (bHLH) type TF, which contain two additional PAS domains and whose efficient DNA binding requires dimerization with another bHLH protein. Since the response element of HIF1A-HIF1B heterodimer (GRACGTGC; HIF1A response element: ACGTGC; JASPAR database http://jaspar.genereg.net/ 30) is very similar to the canonical palindrome consensus sequence called E-box (CACGTG) of bHLH TFs, this opens up the possibility of fine-tuned regulation of *UCP1* gene expression by the Myc/Max/Mad network, Hey2/Hes1, TCFL5, NPAS2 and bHLHE40-41. In addition, ID proteins (Inhibitor of Differentiation 1-3), which appeared in the expanded white gene set and show significantly increased expression in our *FTO* obesity-risk brown adipocyte samples, may be closely related to and involved in these regulatory processes by blocking the DNA binding and transactivation function of bHLH TFs (31) and thus hindering browning. An SNP identified in the HIF1A response element region (rs18006600) may also affect the binding probability of TFs and the regulation of *UCP1* gene expression (Figure 4B; ChIPSummitDB). According to the expression profile of these overlapping TFs (Figure 4F, Upper part), which have the potential to prevent HIF1A-HIF1B binding most of them could repress of the *UCP1* gene-expression, as they show higher expression in our *FTO* obesity-risk brown adipocyte samples, and only *bHLHE40 (DEC1)* and *bHLHE41* appear as possible positive regulators according to their increased expression in our *FTO* normal genotype samples. The other transcriptional factors nearby to the HIF1A-HIF1B heterodimer could cooperate with it to form a complex and/or interact with other regulators and cofactors in the *UCP1* promoter. Their relative expression profile suggests that most of them also may suppress *UCP1* expression, as they are more expressed in *FTO* obesity-risk brown adipocyte samples, but the RXRA-Nr1h3(LXRA) heterodimer and the znf740 could act as activators as their expression is higher in the *FTO* normal genotype samples (Figure 4G; Lower part).

**Figure 4.**
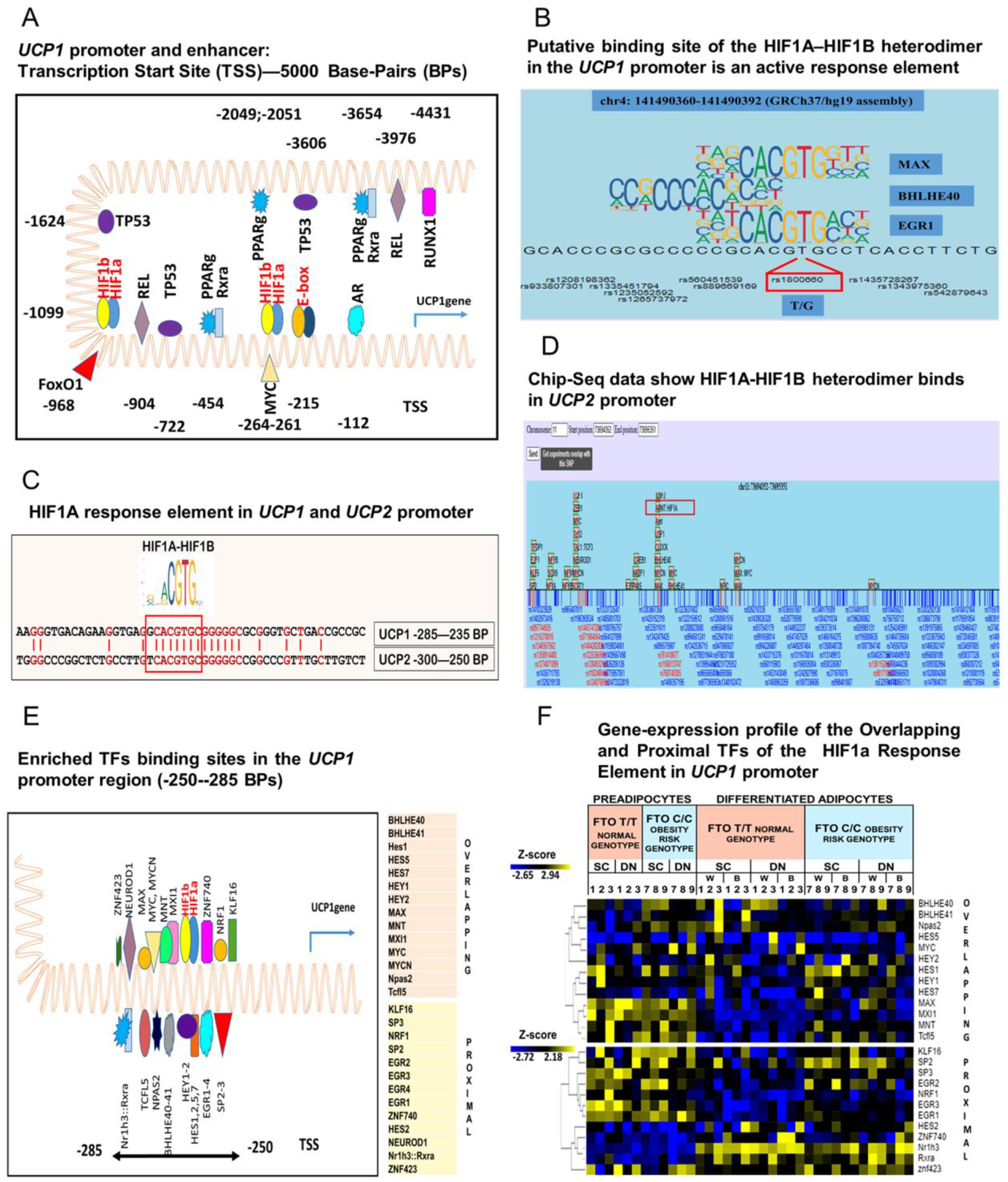
Promoter region of the *UCP1* and *UCP2* genes with enriched TF binding sequence and the gene-expression profile of these TFs. **(A)** The *UCP1* promoter and enhancer region from Transcription Start Site (TSS) to −5000 BasePairs (BPs); TF binding site represent nucleotide position relative to TSS; figure only shows TFs from the 30 core-module proteins **(B)** The chip-seq. data show TFs that bind to the identified HIFA response elements as well as in the immediate vicinity in the *UCP1* promoter and also show the SNPs coinciding with TFs binding sites. The figure was generated with the ChIPSummitDB online software. **(C)** Nucleotide sequence in the homologous promoter region (−235--300 BPs) of the *UCP1* and *UCP2* genes (EPD database: *UCP1, UCP2.2*, Strand [-]), highlighting the similar region (red) and the HIF1A-HIF1B RE in red box (EPD database). **(D)** The schematic figure shows TFs that bind in the promoter region (TSS - 1000 BPs) of the *UCP2* gene and also show the SNPs coinciding with TFs binding sites. The figure was created with the ChIPSummitDB online software. The chromosome region is given according to the hg19 (GRCh37) human reference genome. Red-box show HIF1A-ARNT among the binding TFs **(E)** The *UCP1* promoter: −250--285 BPs show overlapping and proximal TFs binding sites with HIF1A-HIF1B response element. **(F)** Gene-expression profile of the TFs that have Overlapping and Proximal binding site with HIF1A Response Element in *UCP1* promoter.

Regulation of HIF1A protein level in the cell is primarily mediated by inhibition of continuous proteasome degradation initiated by the enzyme Prolyl-hydroxylase, thus gene expression data do not provide sufficient information on the level of HIF1A. Therefore, we examined its presence in differentiated human Adipose Stromal Fraction Cells (hASFCs) from abdominal subcutaneous origin (Abdominal-SC), Methods). Based on preliminary data, in the subcutaneous derived hASVFCs, HIF1A is not or very weakly stabilized in the *in vitro* differentiated adipocytes kept at 21% O_2_ level. However, addition of a cAMP analogue used for thermogenic activation (as demonstrated by UCP1 expression) resulted in elevated HIF1A stabilization, similarly to hypoxic conditions (1% O_2_ level; Methods; Suppl. Figure 5). Consequently, the cell state-dependent presence of HIF1A via a broad range of regulated genes and highly dynamic DNA occupancy could influence the thermogenic phenotype formation through fine-tuned spatiotemporal gene expression regulation in matured adipocytes. This finding may indicate that the regulatory modules and elements revealed by network analysis can generate testable hypothesis.

## DISCUSSION

Exploring a broad database can blur the variation resulting from the specific experimental systems or the sample origin highlighting the common biological differences. In the case of the two meta-analyzes, BATLAS (14) and ProFAT (15), well usable marker-gene sets for determining the browning capacity of the fat cells or tissues were identified. Based on the similarity of the pathways defined by the marker-genes in the two gene sets, we thought that it was worth generating an expanded gene set for deeper analysis of thermogenesis regulation in adipocytes.

Analysis of the expanded gene set makes it clear that the direction of the regulatory processes of gene expression can be differently interpreted if comparative data from preadipocytes are also included. The differentiation process appears to induce the expression of genes which are highly expressed in the thermogenic phenotype, also in the less thermogenic samples. It means that these genes are not suppressed at the chromatin level even in obesity-risk genotype brown adipocytes which can ensure the readiness of immediate heat production by proper stimulation. These genes may be positive modulators of the browning process; however, by itself their expression in a white adipocyte may not be able to turn on thermogenesis. Several recent studies suggest that alterations of the epigenome can influence adipose tissue plasticity in mice and humans (32-37). These studies collectively imply that it is primarily the repression of genes/pathways that potentially trigger transcriptional reprogramming of adipocytes which is in line with our conclusion drawn from the relative expression profile of expanded gene lists when preadipocytes were considered too. Suppression of certain brown or white pathway genes (indirect regulation of thermogenesis by getting rid of interfering proteins) may result in more decisive physiological consequences in contrast to differentially upregulating processes. For example, a recent study using single nucleus RNA sequencing was able to identify a rare population of cells in brown and white adipose tissues that express the enzyme that negatively regulates thermogenesis, and it suggests that downregulation of ALDH1A1 is required for heat-producing adipocyte phenotype (38). Taken together, most of the well-known brown marker-genes that seems to be required for the thermogenic brown adipocyte state may be essential for proper differentiation too, although, at a moderate level, which fits well with the previously demonstrated phenomenon that the cafeteria diet induces adipocyte thermogenesis in addition to differentiation (39)Their amount may affect browning capacity and promote hyperplasia, while a bit paradoxically, the thermogenic state may be induced through broad gene repression processes.

By network analyses we can explore the biological processes, signaling pathways and key proteins that may become important in thermogenesis from a different perspective. Growing evidence indicates that mouse and adult human BAT (Brown Adipose Tissue) have high heterogeneity (8,38,40) and plasticity (41-43) as a response to external cues that makes characterization a challenge, and may give contradictory results (10,11). Generally, the presence of proteins and their interaction in cells is highly dynamic and with the expanded gene set we can explore all possible interactions of the protein network and identify molecular signatures and pathways that potentially play a role in thermogenesis, but otherwise conditional dependent and therefore hard to catch. It could lead to more generally applicable conclusions than single-state studies. In this analyses, the genes belonging to the pathways that shape the brown and white phenotypes do not show one single integrated network, but rather appeared in discrete clusters, which raises the possibility that different processes may form these phenotypes, and we need to know more about their transient states. Nevertheless, this network arrangement suggests that changing phenotype might need action from two directions, so that in addition to activating the browning processes, repression of molecular elements of the white phenotype may also be necessary to generate the desired thermogenesis without pathological consequences. By inducing or inhibiting the expression of proteins occupying a key position in the network or of a cluster, we may be able to initiate more diversified processes that result in the intended functional change. Among the identified central pathways and proteins, the role of the PI3K/PTEN/Akt/FoxO in the regulation of cellular energy metabolism and the thermogenesis of adipocytes has been highly investigated (44,45) and it is well established that the proteasomal degradation of the PTEN protein leads to Akt activation and inhibited, FoxO1-dependent *UCP1* expression in BAT (46). Consistently, mice overexpressing PTEN are protected from metabolic damage, thus PTEN positively regulates energy expenditure and brown adipose function (47).

The role of HIF signaling in the regulation of adipocyte differentiation (48) and maintaining metabolic homeostasis under O_2_-deficient conditions has also been extensively investigated and explored (26,49,01). In obese people, it has been shown that parts of adipose tissue can become acutely or chronically hypoxic (51), which can contribute to the onset and progression of obesity-associated diseases (52,53). Besides this, many studies suggest that HIF1A/HIF2A/HIF3A may play a role in regulating adipocyte differentiation and thermogenesis in even a non-hypoxic environment (24,54-57), however, the molecular details have not been clarified. Tissue-specific HIF1A I.1 expression and protein accumulation has been reported at the early stage of adipogenesis in normoxic condition (58), and it has also been demonstrated that norepinephrine without UCP1 expression can stabilise HIF1A in brown adipocytes (25). However, the O_2_ concentration has not been studied at the cellular level so we cannot know for sure whether the cells were actually in a state of normoxia or hypoxia. In addition, in a Genome-Wide Association Study, explored of a European-descent individuals’ data, the *HIF1AN* rs17094222 loci was identified as a Body Mass Index (BMI) associated loci (59) and in a recent large-scale epigenome-wide association study it was found that DNA methylation at the *HIF3A* site was associated with BMI (610HIF1A was among the recruited proteins to the *UCP1 cis*-regulatory elements in response to the newly identified browning regulators, the FGF6/FGF9 stimulation in murine brown adipocytes (61). Recently Basse et al. showed that HIF1A expression is important for basal and beta-adrenergic stimulated (by isoproterenol) expression of glycolytic enzymes and necessary for maximum glucose metabolism in thermogenic adipocytes reducing the intense mitochondrial function generated ROS (Reactive Oxygen Species) and its damaging effect (26), contrary to study which demonstrated that adipose HIF1A overexpression inhibits thermogenesis and cellular respiration in brown adipose tissue, promoting obesity in the setting of reduced ambient temperature (57). Notwithstanding, although its role in shaping the thermogenic adipocyte phenotype remains to be clarified, the hitherto unexplored possibility that HIF1A directly binds to the promoter region and regulates *UCP1* gene-expression may shed new light on the regulation of thermogenicity of adipocytes and reveal promising targets for the pharmacological treatment of obesity.

## METHODES

### Ethics statement

Tissue collection was carried out with the guidelines of the Helsinki Declaration and was approved by the Medical Research Council of Hungary (20571-2/2017/EKU) followed by the EU Member States’ Directive 2004/23/EC on presumed consent practice for tissue collection. Prior to the surgical procedure, written consent was obtained from all participants.

### Isolation, Cell Culture, Differentiation and Treatments of hASC and Simpson-Golabi-Behmel Syndrome (SGBS) Cells

The hASCs were isolated from Subcutaneous (SC) adipose tissue. The volunteer underwent a planned surgical treatment, 48 years with BMI< 19.7. Patients were known without malignant tumor, diabetes or with abnormal thyroid hormone levels at the time of surgery. hASCs were isolated and cultivated as it is already described in Tóth et al (18). The absence of mycoplasma was checked by polymerase chain reaction (PCR) analysis (PCR Mycoplasma Test Kit I/C, PromoCell cat# PK-CA91). The SC abdominal adipocytes from hASCs were isolated and grown till confluence state, than differentiated with white differentiation cocktail on 6 well plates at 37 °C in 5% CO_2_, with white cocktail, according to previously described protocols (18). Briefly, cells were grown in DMEM-F12-HAM (Sigma-Aldrich cat#D8437) medium supplemented with 10% FBS (Gibco cat#10270106); 10% EGM2 (Lonza #CC3162), 1% Biotin (Sigma-Aldrich cat#B4639), 1% Pantothenic acid (Sigma-Aldrich cat#P5155), 1% Streptomycin-Penicillin (Sigma-Aldrich cat#P4333), white differentiations were induced at the first 4 days by hormonal cocktails in serum and additive-free DMEM-F12-HAM (Sigma-Aldrich cat#D8437) medium that contain biotin, pantothenic acid, apo-transferrin (Sigma-Aldrich cat#T2252), insulin (Sigma-Aldrich cat#I9278), T3 (Sigma-Aldrich cat#T6397), dexamethasone (Sigma-Aldrich cat#D1756), hydrocortisone (Sigma-Aldrich cat#H0888) rosiglitazone (Cayman Chemicals cat#71740) and IBMX (Sigma-Aldrich cat#I5879). Later, for 10 days rosiglitazone, dexamethasone and IBMX were omitted from the media. To compose the brown media for termogenic induction serum and additive-free DMEM-F12-HAM (Sigma-Aldrich cat#D8437) medium that contain biotin, pantothenic acid, apo-transferrin (Sigma-Aldrich cat#T2252), insulin (Sigma-Aldrich cat#I9278), T3 (Sigma-Aldrich cat#T6397) and rosiglitason (Cayman Chemicals cat#71740) were present in the cocktail, and the insulin was present at 40x higher concentration than in the white differentiation cocktail.

### Thermogenic activation and Hypoxic Experiment

Differentiated adipocytes on 6 well plates were used for the experiment. Two treatments were performed in parallel (3 wells/treatment): Thermogenic cocktail (Brown media supplemented with 500 μM cAMP analog (dibutyril-cAMP; Sigma-Aldrich cat#D0627) (9)) was applied for 16 h to induce thermogenesis, or hypoxic condition (1% O_2_) was maintained for 16 h in hypoxic chamber, while the control group was the untreated differentiated adipocytes. The hypoxic gas-mix contains 1% O_2_, 5% CO_2_, 94% N_2_.

### Western blotting

The cellular lysate was obtained using 2X denaturing buffer (100ml: 25ml TRIS pH 6.8; 45ml MilliQ Water; 20ml glycerol; 10ml MEA; 4g SDS, Bromophenol Blue) diluted to 1:1 with urea (8M). Briefly, 20μl protein from cell culture lysates was separated using 10% acrylamide gel, transferred to PVDF Immobilon-P Transfer Membrane (Merck-Millipore, Darmstadt, Germany), blocked with 5% milk powder in TTBS (0.1% Tween 20 in TBS), and incubated overnight at 4 °C separately with UCP1 (1:750 dilution in 1% milk powder in TTBS; R&D Systems, Minneapolis, MN, USA, MAB6158), HIF1A (1:750 dilution in 1% milk powder in TTBS; BD cat#610958) and beta-Tubulin (1:10000 dilution in 1% milk powder in TTBS Sigma-Aldrich cat#F2168) antibodies. Following overnight incubation, blots were incubated with HRP-conjugated secondary anti-mouse antibody for 1h (1:5000 dilutions in 1% milk powder in TTBS; 1:5000, Advansta, R-05071-500) and exposed to x-ray films using chemiluminescent detection method (ECL, Merck-Millipore, MA, USA).

### Generation of the Expanded gene set

BATLAS and ProFAT brown and white marker-genes (basic marker-genes) were subjected to Gene Set Enrichment Analysis (GSEA) separately (17). We explored the two marker-gene lists to identify enrichment of specific Functional Gene Sets that is annotated to a certain pathway. Enriched pathways are pathways that annotated genes of a pathway are represented in the gene sets significantly higher than by chance. The enriched pathways were identified by the open source The Kyoto Encyclopedia of Genes and Genomes (KEGG) and REACTOME pathway analysers which are integrated into the STRING computer tool (software) (https://string-db.org; 19). Enriched pathway: annotated genes of the pathway are represented in the two gene-sets significantly higher than the random probability. Based on the KEGG (https://www.genome.jp/kegg/pathway.html as of February 2020) (19) and REACTOME (http://www.reactome.org as of February 2020) (20) databases we retrieve annotated gene sets for enriched pathways. For generating the expanded brown gene set, only genes belonging to those pathways that were enriched for both BATLAS and ProFAT were collected (Figure 1C, D, Table 1), in the case of the basic brown marker-genes. In the case of the basic white marker-genes, only the BATLAS genes enriched pathways were used, because, in the case of ProFAT, the 6 white genes did not define any enriched pathway (Figure 1C, D). Pathways that are too general and have a large number of genes (more than 600) were excluded from the analysis, thus Metabolism (1250 KEGG genes, 2032 REACTOME genes) and Metabolism of Lipid (721 REACTOME genes) among brown genes and Signal Transduction (2605 REACTOME genes) and Developmental Biology (1023 REACTOME genes) pathways for white genes were excluded from the expanded gene sets (Supplementary Table 4, 5, 6, 7).

### GSEA analyses

Gene Set Enrichment Analyses was performed with STRING computer tool, as Gene Ontology (GO) algorithms and statistical analyses, KEGG and REACTOME pathway analysers are integrated into the STRING computer tool (https://string-db.org; 21).

### Interactome analyses

We used the computational tool STRING (https://string-db.org; 21) for generating protein-based interactomes, where nodes represent a protein and edges represent protein-protein based interactions. STRING and Cytoscape applications were used for the visualisation of interactomes. Exploring the interaction of proteins encoded by the expanded brown and white gene-sets we used the default median interaction confidence score (0.4) as a threshold. The STRING database considers seven specific types of evidence for the interaction of two proteins that contribute to overall interaction confidence score, and we used all of them. Further analyzes of the STRING network data were performed in the Igraph package (62; https://igraph.org.) of the R interactive statistical environment (63) to compute betweenness scores and number of bridges of the interacting nodes. We calculated the betweenness centrality score to identify important genes that have the most central positions in connecting different functional modules and the number of bridges to learn how many modules are connected by this gene. Interaction score confidence values (0.4) were used to visualize the network, as indicated in the figures. Unsupervised MCL clustering were used from the ClusterMaker2 application downloaded in Cytoscape softwere to explore subclusters of the generated Interactome of the 3705 Brown and White Pathways genes encoded proteins. Interaction score confidence value: 0.4. MCL parameters: Granularity parameter: 2; Array source: stringdb: score; Edge wieght cutoff: 80; No restored inter-cluster edges after layout.

### Heatmap visualization

Hierarchical cluster analyses and heat map visualization of the relative gene-expression data was performed in the Morpheus web tool (https://software.broadinstitute.org/morpheus/) using Pearson correlation of rows and complete linkage based on calculated z-score of DESeq normalized and filtered data (low-expressed and outlier) after log2 transformation (18). Z score is calculated as (X - m)/s, where X is the value of the element, m is the mean, and s is the SD. Heatmaps shows Samples 1-3 with *FTO* T/T-risk-free allele (donor 1-3) and samples 7-9 with C/C-obesity-risk alleles (donor 7-9), we left out the *FTO* T/C heterozygous samples (donor 4-6) which are shown for some case in the paper Tóth et al. (18).

### Origin of the Transcriptomic Data

Ethics Statement and Obtained Samples, Isolation, and Differentiation of human Adipose Stromal Cells (hASCs), as well as RNA and DNA Isolation, Genotyping, RNA-Sequencing is already described in Tóth et al.(18). The hASCs were isolated from paired DN (Deep Neck) and SC (Subcutaneous Neck) adipose tissue samples of nine donors, three of each *FTO* rs1421085 genotype: T/T-risk-free, T/C-heterozygous, and C/C-obesity-risk. After cultivation, both brown and white differentiations were induced by hormonal cocktails in serum and additive-free DMEM-F12-HAM medium.

### Transcription Factors binding sites in the UCP1 promoter

The eukaryotic promoter database (EPD; https://epd.epfl.ch) was used to determine whether the Transcription Factor (TF) have binding site (response element) in the *UCP1* promoter region (23). The EPD contains comprehensive organisms-specific transcription start site (TSS) collections automatically derived from next generation sequencing (NGS) data and corresponding promoter region (−5000 and +100 Base pairs from TSS). We use the library of the TF Motifs (JASPAR CORE 2018 vertebrates;30). The consensus sequence of the TFs was identified by the JASPAR (http://jaspar.genereg.net/) database, which is an open-access database of curated, non-redundant TF-binding profiles (30).

Chip-Atlas (http://dbarchive.biosciencedbc.jp; 27) and ChIPSummitDB database (University of Debrecen; http://summit.med.unideb.hu; 28) were used to investigate experimental data about HIF1A and other TFS binding to *UCP1* and *UCP2* promoter. Both online tools collect, organize and visualise the results of the published chip-seq. experiments. The chromosome region is given according the hg19 (GRCh37) human reference genome in the case in ChIPSummitDB, otherwise the GRCh38/hg38 genome assembly was used.

### Comparative Genomic Analyses

For comparative genomic analysis we used the UCSC genome browser (University of California Genomics Institute, Santa Cruz, https://genome-euro.ucsc.edu) and tracked the close vicinity of the HIF1A response element in the *UCP1* promoter. The tracking shows multiple alignments of 100 vertebrate species and measurements of evolutionary conservation.

### Data and Software Availability

The accession number for the RNA-seq data from human studies reported in this paper is deposited to the Sequence Read Archive (SRA) database (https://www.ncbi.nlm.nih.gov/sra) under accession number PRJNA607438.. Algorithms for calculation of the Betweenness Centrality Score and Number of Bridges based on Protein Network Interaction confidence values identified by STRING computational tool was developed in this work and can be accessed at http://github.com/zbartab/brownRNA.

## Supporting information

BATLAS genes

Pro FAT genes

Supplementary Table Title and Legends

## Acknowledgements

We kindly acknowledge the human subcutaneous adipose tissue sample to Zoltán Péter. We thank Erik Czipa for introducing the use of the ChIPSummitDB database; Ifj. Tamás Székely, Bálint László Bálint and Endre Kristóf for reviewing the manuscript and giving useful advice and Jennifer Nagy for excellent technical assistance. This research was funded by the European Union and the European Regional Development Fund (GINOP-2.3.2-15-2016-00006) and we are thankful for this.

## Author Contributions

Conceptualization, B.B.T., methodology, B.B.T. and Z.B.; software, B.B.T. and Z.B.; formal analysis, B.B.T and Z.B.; visualization, B.B.T.; investigation, B.B.T.; data curation, B.B.T.; writing—original draft preparation, B.B.T., and L.F.; writing—review and editing, B.B.T., L.F. and Z.B.; supervision, L.F.; project administration, L.F; funding acquisition, L.F. All authors have read and agreed to the published version of the manuscript.

## Declaration of Interests

The authors declare no conflict of interest.

## Supplemental Tables titles and legends

**Supplementary table 1.**

**List of BATLAS Marker-genes**

**Supplementary table 2.**

**List of ProFAT Marker-genes**

**Supplementary table 3. Significantly enriched pathways based on BATLAS, ProFAT marker-genes, and genes differentially expressed in human neck adipocyte samples based on the presence of the FTO obesity-risk allele. (A) KEGG pathways (B) REACTOM** pathways identified by BATLAS, ProFAT Brown marker-genes and the genes expressed lower in *FTO* C/C (obesity risk) samples. **(C) KEGG pathways (D) REACTOM** pathways identified by BATLAS White marker-genes and the genes expressed Higher in *FTO* C/C samples. Red letters: Common Pathways that enriched based on BATLAS and ProFAT marker-genes; Blue letters: Common Pathways in *FTO* based DEGs and BATLAS or ProFAT; Italic font: Common enriched Pathways in *FTO* based DEGs and BATLAS and ProFAT.

**Supplementary table 4.**

**KEGG pathway analyses of the BATLAS (98 genes) and ProFAT (53 genes) Brown marker-genes.** Working sheets show the annotated genes for the enriched pathways based on KEGG database.

**Supplementary table 5.**

**REACTOME pathway analyses of the BATLAS (98 genes) and ProFAT (53 genes) Brown marker-genes.** Working sheets show the annotated genes for the enriched pathways based on REACTOME database.

**Supplementary table 6.**

**KEGG pathway analyses of the BATLAS white marker-genes (21 genes).** Working sheets show the annotated genes for the enriched pathways based on KEGG database.

**Supplementary table 7.**

**REACTOME pathway analyses of the BATLAS white marker-genes (21 genes).** Working sheets show the annotated genes for the enriched pathways based on REACTOME database.

**Supplementary table 8.**

**List of the genes belongs to the downregulated gene cluster (dark red box; 175 genes in Figure 2 B) identified by the hierarchical cluster analyses of the gene-expression data of the expanded Brown gene-set and their GSEA.**

## Supplementary Figures

**Supplementary Figure 1.**
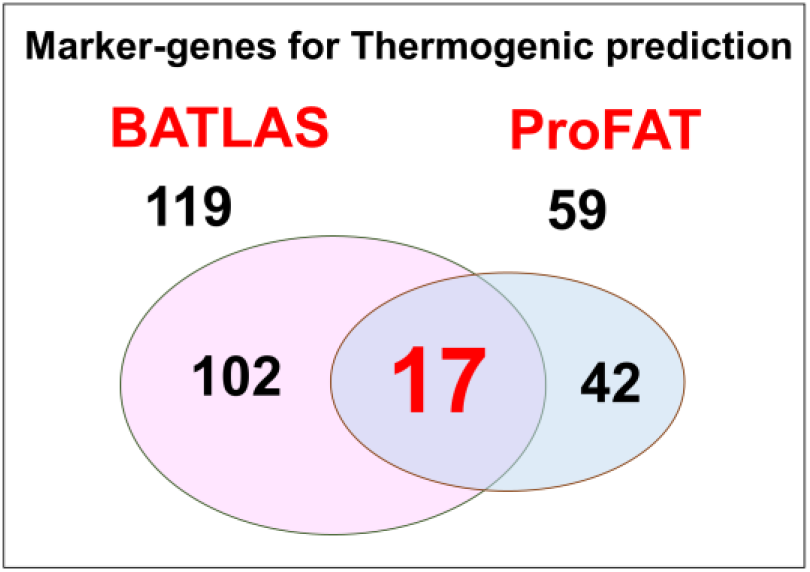
Marker-genes of BATLAS and ProFAT. Venn diagrams show the number of marker-genes from BATLAS and ProFAT databases

**Supplementary Figure 2.**
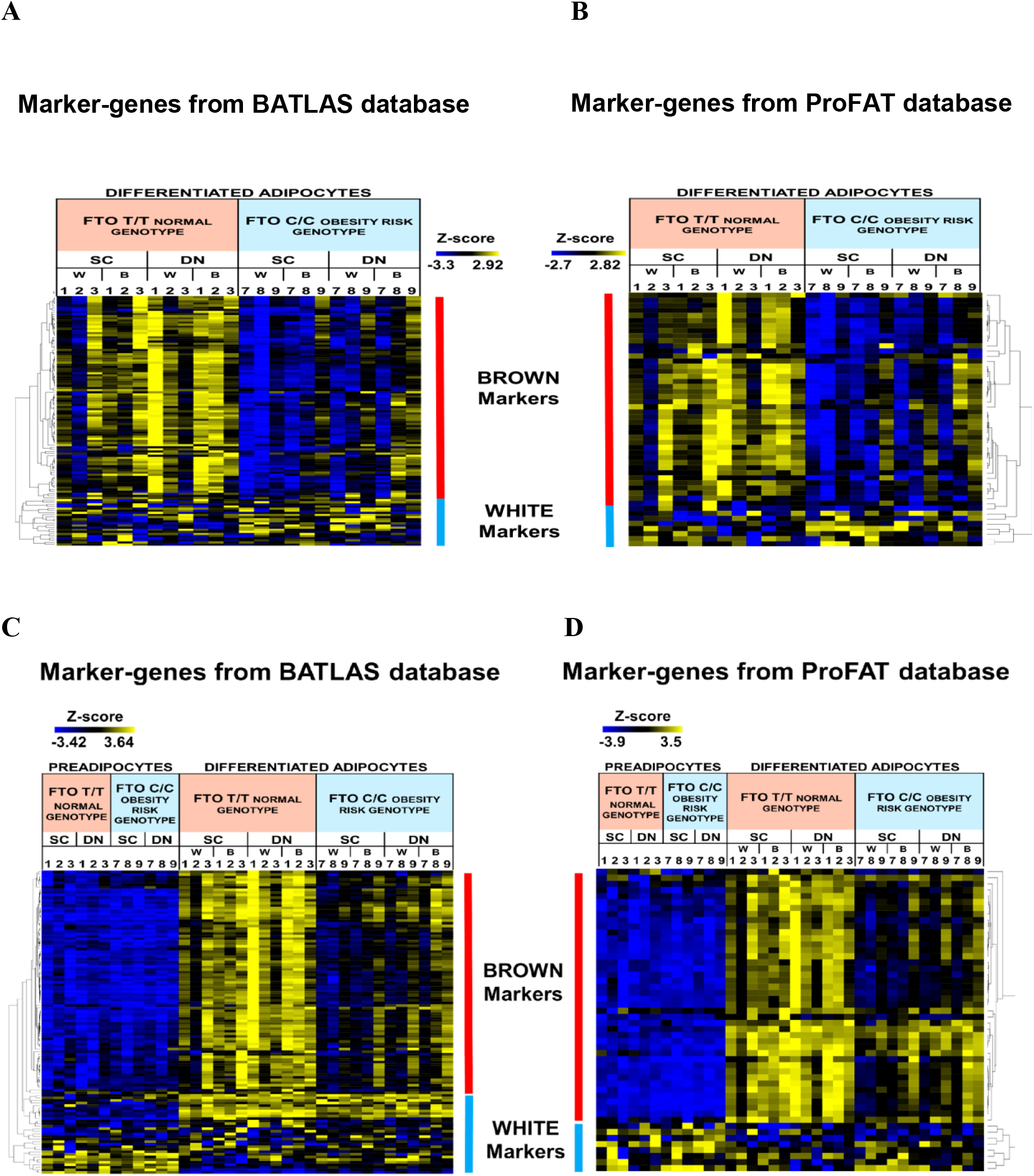
Relative gene-expression profile of marker-genes from BATLAS and ProFAT. Relative gene-expression profile of marker-genes from BATLAS **(A)** and ProFAT **(B)** based on the presence of FTO obesity-risk alleles in differentiated Subcutaneous and Deep-neck adipocytes *n* = 6; samples1-3 with *FTO* T/T obesity-risk-free allele (donor 1-3) and samples 7-9 with C/C-obesity-risk alleles (donor 7-9) the *FTO* T/C heterozygous samples are not shown (donor 4-6) which was presented in some case in the paper Tóth et al., 2020. (BATLAS **(C)** and ProFAT **(D)** marker-genes expression profile based on the presence of FTO obesity-risk alleles in pre and differentiated Subcutaneous and Deep-neck adipocytes *n* = 6

**Supplementary Figure 3.**
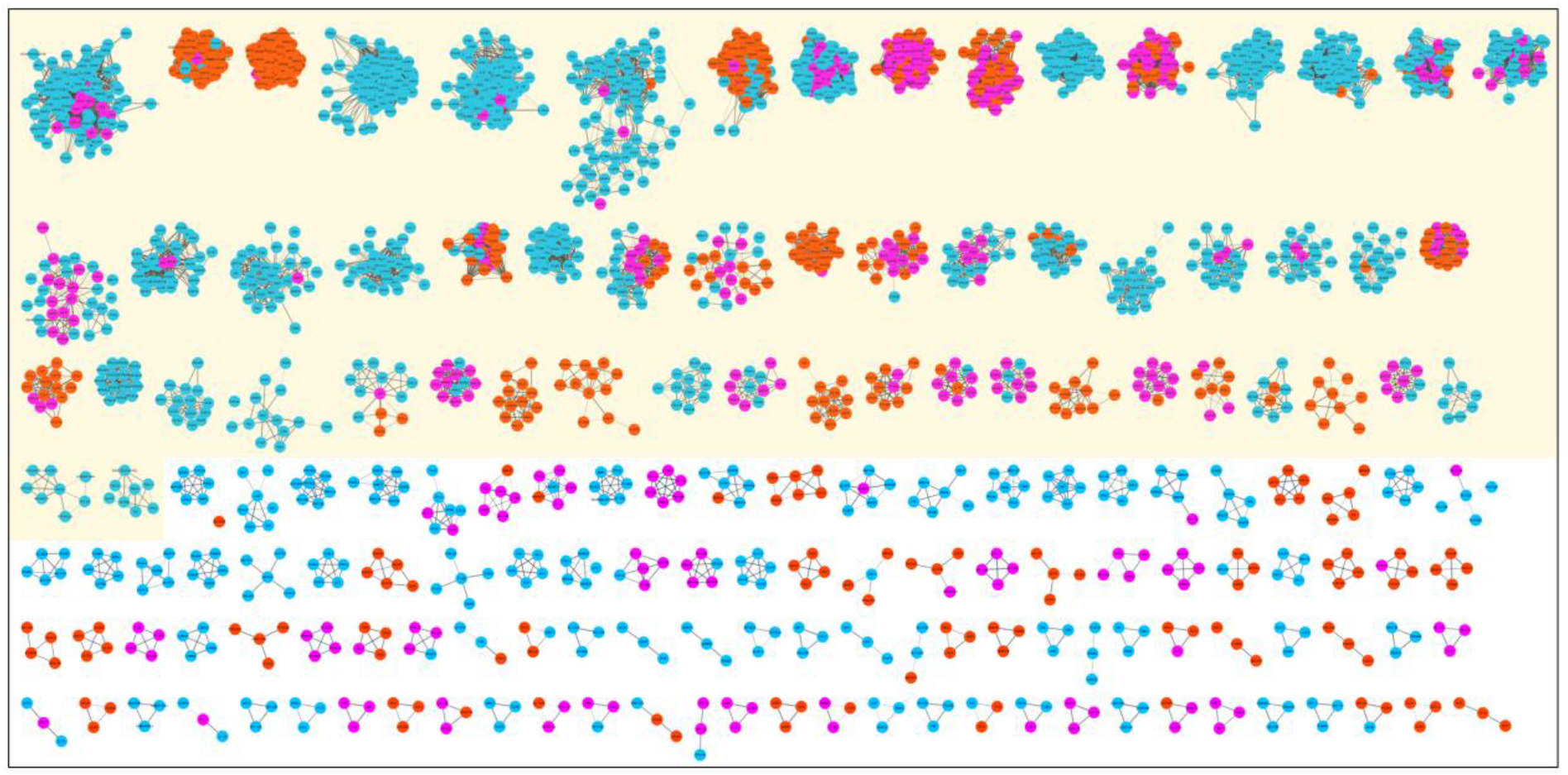
Unsupervised MCL of the Interactome network of the 3705 Brown and White Pathways genes encoded proteins generated sub-clusters (Cytoscape); Nodes represent proteins; edges represent protein-protein interactions. Figure only show clusters contains more than 3 proteins. Cream colour highlights clusters contains 8 or more proteins. Red nodes: Brown pathway proteins; Blue nodes: White pathway proteins; Magenta: proteins appeared in both Brown and White Pathway

**Supplementary Figure 4.**
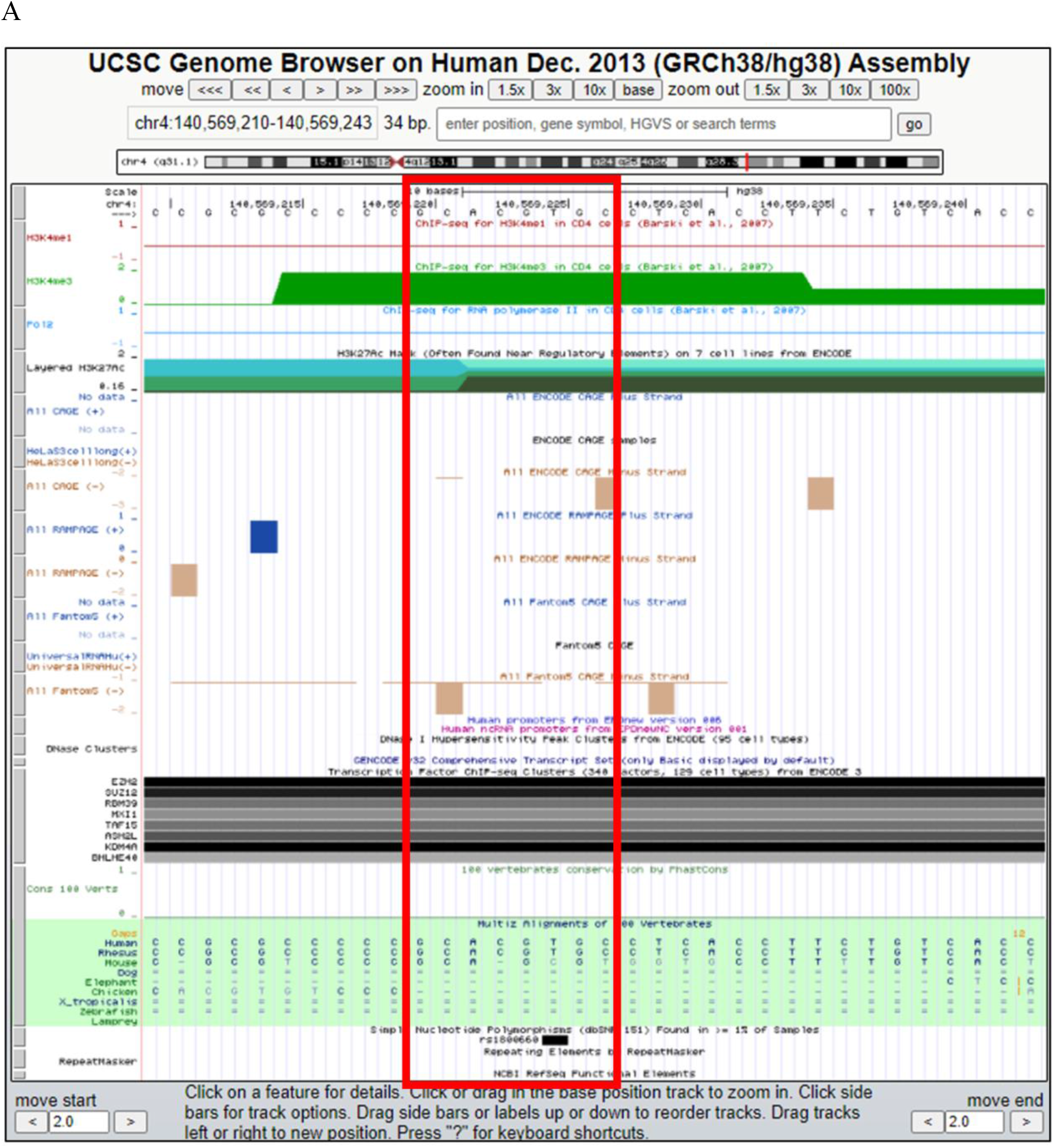
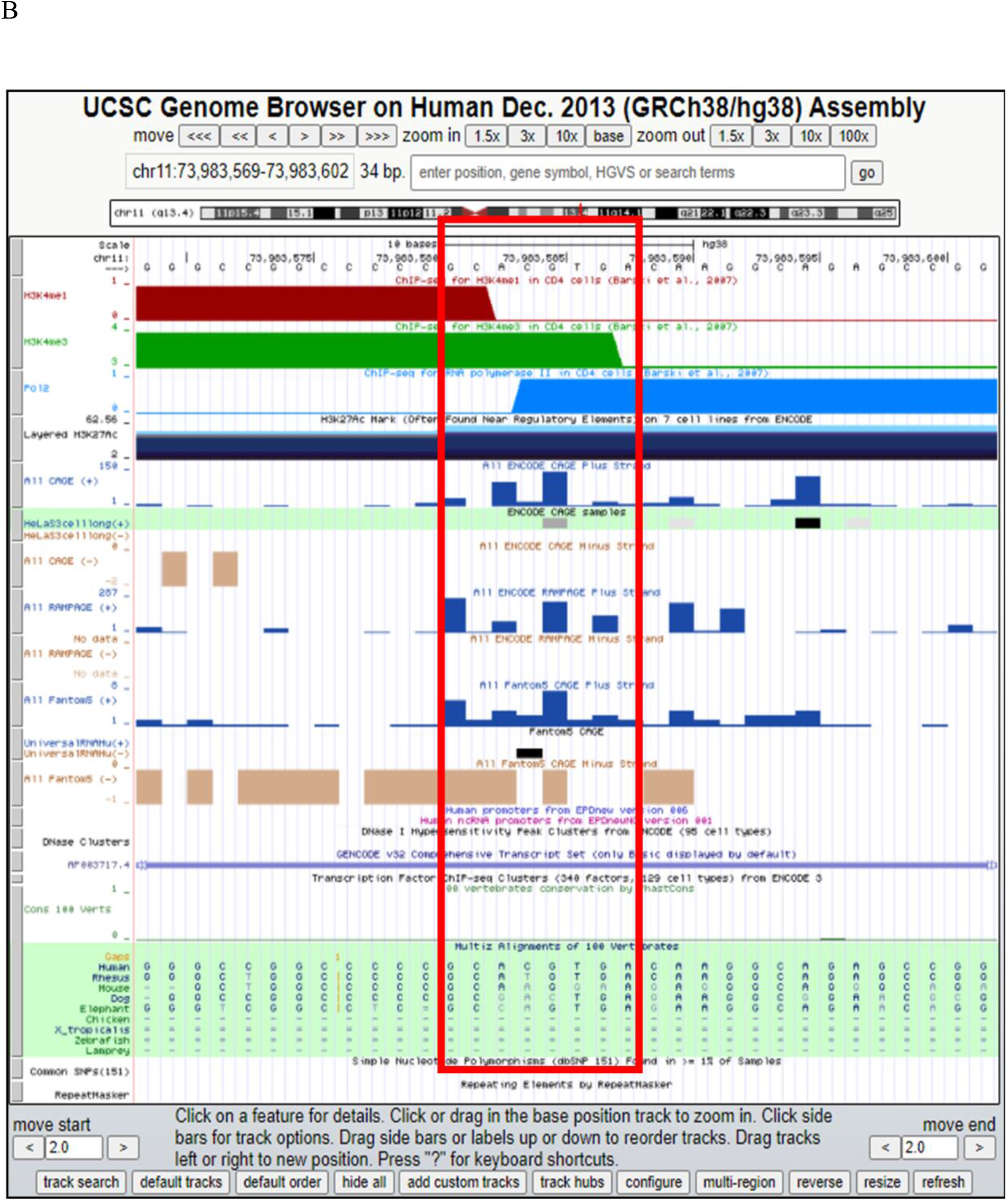
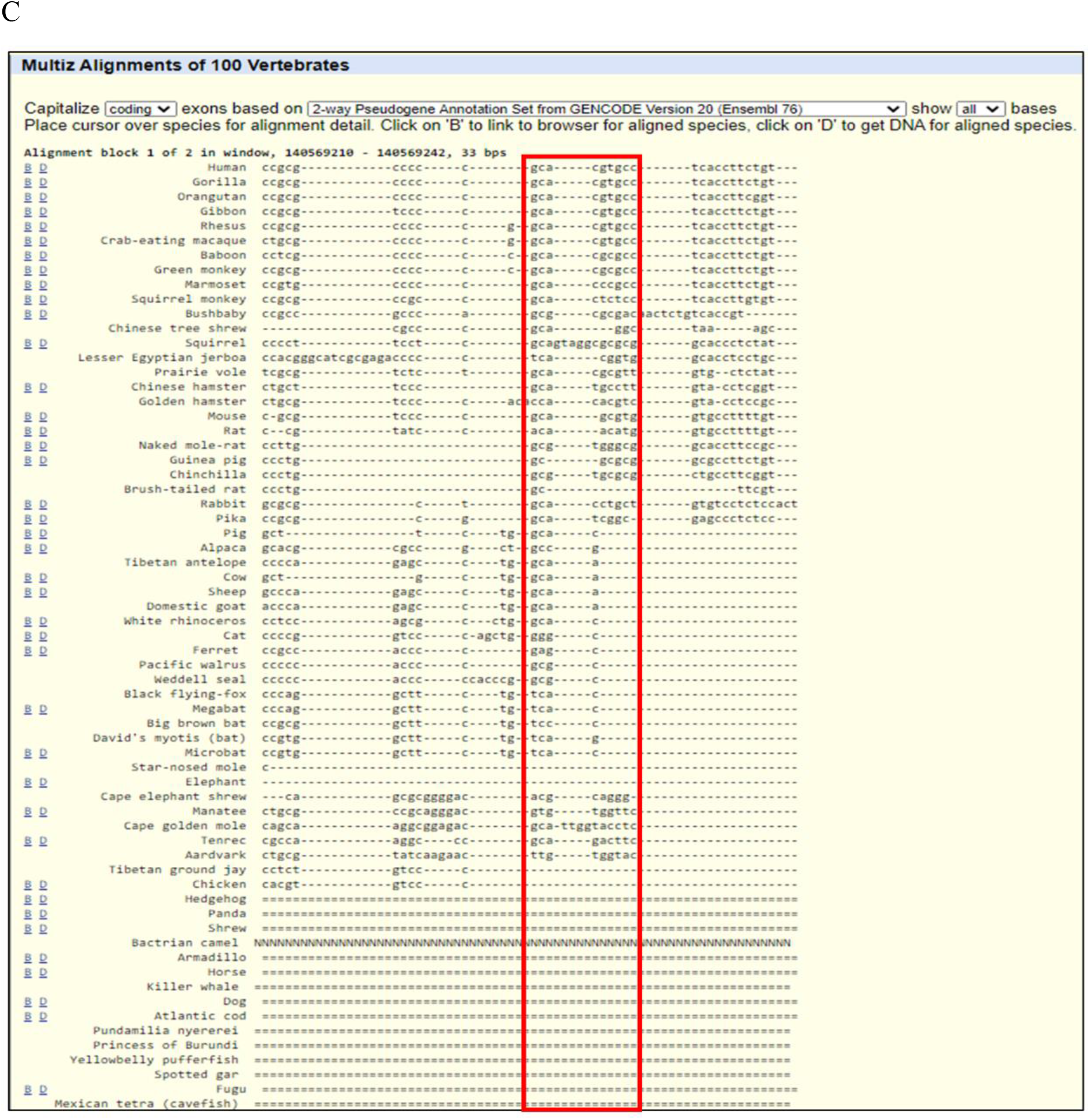
Comparative genomic analysis by UCSC genome browser. **(A)** UCSC genome browser overview of the *UCP1* promoter where HIF1A binding sequence is enriched (red box) with Histone methylation state and alignments of orthologues sequence from other species highlighted in green; Strand [+] (https://genome-euro.ucsc.edu/; EPD Viewer HUB). **(B)** UCSC genome browser overview of the *UCP2* promoter where HIF1A binding sequence is enriched (red box) with Histone methylation state and alignments of orthologues sequence from other species highlighted in green; Strand [+] (https://genome-euro.ucsc.edu/; EPD Viewer HUB). **(C**) Multiz Alignments of 100 Vertebrates data show the detailed sequence homology of the different species in the *UCP1* promoter where HIF1A binding sequence is enriched (red box) (Further species with no homologous sequence did not shown (https://genome-euro.ucsc.edu/cgi-bin/hgc?c=chr4&l=140569209&r=140569243&o=140569209&t=140569243&g=multiz100way&i=multiz100way&db=hg38).

**Supplementary Figure 5.**
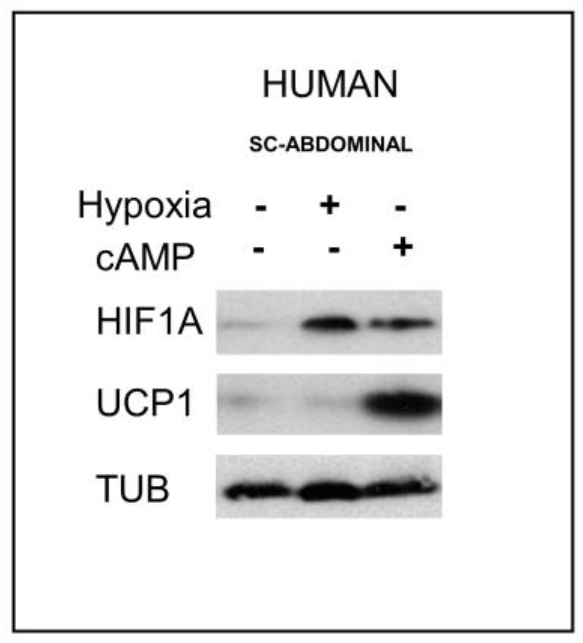
Effect of thermogenic induction and Hypoxic condition on the protein level of HIF1A and UCP1. Immunoblots show HIF1A and UCP1 protein levels in cAMP analog-induced thermogenesis (16h) and hypoxic environment (16h) in differentiated adipocyte sample derived from human subcutaneous stromal vascular fraction, using beta tubulin (TUB) as a loading control. SC: Subcutaneous;

## Notes

https://data.mendeley.com/datasets/hsj8wsfz7t/draft?a=fe332df2-48c7-42ed-b6aa-b635da0c6cac

### Competing Interest Statement

The authors have declared no competing interest.

### Summary of Updates

Figure 5. has been modified and relocated to supplement files as Supplement Figure 5.

