## Supplementary material for "Regulatory Modules of Human Thermogenic Adipocytes: Functional Genomics of Meta-Analyses Derived Marker-Genes": BATLAS genes

Table 1. BATLAS Marker-gene list.

| **BATLAS Gene ID** | Mouse Code | Human Code | Adipose Tissue marker type |
| --- | --- | --- | --- |
| **ACSL5** | ENSMUSG00000024981 | ENSG00000197142 | brown |
| **MT-ND3** | ENSMUSG00000064360 | ENSG00000198840 | brown |
| **PANK1** | ENSMUSG00000033610 | ENSG00000152782 | brown |
| **UCP1** | ENSMUSG00000031710 | ENSG00000109424 | brown |
| **MT-CO3** | ENSMUSG00000064358 | ENSG00000198938 | brown |
| **LETMD1** | ENSMUSG00000037353 | ENSG00000050426 | brown |
| **GK** | ENSMUSG00000025059 | ENSG00000198814 | brown |
| **GOT1** | ENSMUSG00000025190 | ENSG00000120053 | brown |
| **CPT1B** | ENSMUSG00000078937 | ENSG00000205560 | brown |
| **KCNK3** | ENSMUSG00000049265 | ENSG00000171303 | brown |
| **MT-ND2** | ENSMUSG00000064345 | ENSG00000198763 | brown |
| **ADCY3** | ENSMUSG00000020654 | ENSG00000138031 | brown |
| **NDUFAB1** | ENSMUSG00000030869 | ENSG00000004779 | brown |
| **DES** | ENSMUSG00000026208 | ENSG00000175084 | brown |
| **BSG** | ENSMUSG00000023175 | ENSG00000172270 | brown |
| **PPIF** | ENSMUSG00000021868 | ENSG00000108179 | brown |
| **COX5A** | ENSMUSG00000000088 | ENSG00000178741 | brown |
| **SUCLG1** | ENSMUSG00000052738 | ENSG00000163541 | brown |
| **CYC1** | ENSMUSG00000022551 | ENSG00000179091 | brown |
| **PPARGC1B** | ENSMUSG00000033871 | ENSG00000155846 | brown |
| **MRPL34** | ENSMUSG00000034880 | ENSG00000130312 | brown |
| **AMACR** | ENSMUSG00000022244 | ENSG00000242110 | brown |
| **NDUFS8** | ENSMUSG00000059734 | ENSG00000110717 | brown |
| **NDUFB2** | ENSMUSG00000002416 | ENSG00000090266 | brown |
| **CIAPIN1** | ENSMUSG00000031781 | ENSG00000005194 | brown |
| **SOD2** | ENSMUSG00000006818 | ENSG00000112096 | brown |
| **HADHA** | ENSMUSG00000025745 | ENSG00000084754 | brown |
| **UQCRC1** | ENSMUSG00000025651 | ENSG00000010256 | brown |
| **ACADVL** | ENSMUSG00000018574 | ENSG00000072778 | brown |
| **DLST** | ENSMUSG00000004789 | ENSG00000119689 | brown |
| **HCCS** | ENSMUSG00000031352 | ENSG00000004961 | brown |
| **CRLS1** | ENSMUSG00000027357 | ENSG00000088766 | brown |
| **IDH3B** | ENSMUSG00000027406 | ENSG00000101365 | brown |
| **COX10** | ENSMUSG00000042148 | ENSG00000006695 | brown |
| **DNAJA3** | ENSMUSG00000004069 | ENSG00000103423 | brown |
| **ETFA** | ENSMUSG00000032314 | ENSG00000140374 | brown |
| **NDUFB7** | ENSMUSG00000033938 | ENSG00000099795 | brown |
| **NDUFS3** | ENSMUSG00000005510 | ENSG00000213619 | brown |
| **CHCHD3** | ENSMUSG00000053768 | ENSG00000106554 | brown |
| **SLC25A39** | ENSMUSG00000018677 | ENSG00000013306 | brown |
| **IDH3A** | ENSMUSG00000032279 | ENSG00000166411 | brown |
| **COQ6** | ENSMUSG00000021235 | ENSG00000119723 | brown |
| **COX7A2** | ENSMUSG00000032330 | ENSG00000112695 | brown |
| **OGDH** | ENSMUSG00000020456 | ENSG00000105953 | brown |
| **VWA8** | ENSMUSG00000058997 | ENSG00000102763 | brown |
| **AGPAT3** | ENSMUSG00000001211 | ENSG00000160216 | brown |
| **MTIF2** | ENSMUSG00000020459 | ENSG00000085760 | brown |
| **UQCR10** | ENSMUSG00000059534 | ENSG00000184076 | brown |
| **NDUFB9** | ENSMUSG00000022354 | ENSG00000147684 | brown |
| **SDHC** | ENSMUSG00000058076 | ENSG00000143252 | brown |
| **ACO2** | ENSMUSG00000022477 | ENSG00000100412 | brown |
| **NDUFS2** | ENSMUSG00000013593 | ENSG00000158864 | brown |
| **ECH1** | ENSMUSG00000053898 | ENSG00000104823 | brown |
| **DNAJC11** | ENSMUSG00000039768 | ENSG00000007923 | brown |
| **SDC4** | ENSMUSG00000017009 | ENSG00000124145 | brown |
| **PHB2** | ENSMUSG00000004264 | ENSG00000215021 | brown |
| **SGPL1** | ENSMUSG00000020097 | ENSG00000166224 | brown |
| **MRPL15** | ENSMUSG00000033845 | ENSG00000137547 | brown |
| **OXNAD1** | ENSMUSG00000021906 | ENSG00000154814 | brown |
| **UQCC1** | ENSMUSG00000005882 | ENSG00000101019 | brown |
| **UQCRB** | ENSMUSG00000021520 | ENSG00000156467 | brown |
| **TIMM44** | ENSMUSG00000002949 | ENSG00000104980 | brown |
| **HSPD1** | ENSMUSG00000025980 | ENSG00000144381 | brown |
| **NDUFA13** | ENSMUSG00000036199 | ENSG00000186010 | brown |
| **TBRG4** | ENSMUSG00000000384 | ENSG00000136270 | brown |
| **POLN** | ENSMUSG00000045102 | ENSG00000130997 | brown |
| **ECSIT** | ENSMUSG00000066839 | ENSG00000130159 | brown |
| **DLD** | ENSMUSG00000020664 | ENSG00000091140 | brown |
| **NDUFV1** | ENSMUSG00000037916 | ENSG00000167792 | brown |
| **AKAP1** | ENSMUSG00000018428 | ENSG00000121057 | brown |
| **GLRX5** | ENSMUSG00000021102 | ENSG00000182512 | brown |
| **NDUFS1** | ENSMUSG00000025968 | ENSG00000023228 | brown |
| **THEM4** | ENSMUSG00000028145 | ENSG00000159445 | brown |
| **SLC25A11** | ENSMUSG00000014606 | ENSG00000108528 | brown |
| **MRPS7** | ENSMUSG00000046756 | ENSG00000125445 | brown |
| **POLDIP2** | ENSMUSG00000001100 | ENSG00000004142 | brown |
| **PDHA1** | ENSMUSG00000031299 | ENSG00000131828 | brown |
| **SDHA** | ENSMUSG00000021577 | ENSG00000073578 | brown |
| **BCKDHB** | ENSMUSG00000032263 | ENSG00000083123 | brown |
| **HSPA9** | ENSMUSG00000024359 | ENSG00000113013 | brown |
| **GATB** | ENSMUSG00000028085 | ENSG00000059691 | brown |
| **EHHADH** | ENSMUSG00000022853 | ENSG00000113790 | brown |
| **IMMT** | ENSMUSG00000052337 | ENSG00000132305 | brown |
| **PTCD3** | ENSMUSG00000063884 | ENSG00000132300 | brown |
| **MRPS18B** | ENSMUSG00000024436 | ENSG00000204568 | brown |
| **NDUFAF5** | ENSMUSG00000027384 | ENSG00000101247 | brown |
| **MARCH5** | ENSMUSG00000023307 | ENSG00000198060 | brown |
| **ACADS** | ENSMUSG00000029545 | ENSG00000122971 | brown |
| **MRPS5** | ENSMUSG00000027374 | ENSG00000144029 | brown |
| **MRPS22** | ENSMUSG00000032459 | ENSG00000175110 | brown |
| **ECHS1** | ENSMUSG00000025465 | ENSG00000127884 | brown |
| **PCK1** | ENSMUSG00000027513 | ENSG00000124253 | brown |
| **ACSF2** | ENSMUSG00000076435 | ENSG00000167107 | brown |
| **TIMM50** | ENSMUSG00000003438 | ENSG00000105197 | brown |
| **C1QBP** | ENSMUSG00000018446 | ENSG00000108561 | brown |
| **FLAD1** | ENSMUSG00000042642 | ENSG00000160688 | brown |
| **AURKAIP1** | ENSMUSG00000065990 | ENSG00000175756 | brown |
| **ACAT1** | ENSMUSG00000032047 | ENSG00000075239 | brown |
| **NNAT** | ENSMUSG00000067786 | ENSG00000053438 | white |
| **PRKCDBP** | ENSMUSG00000037060 | ENSG00000170955 | white |
| **IGF1** | ENSMUSG00000020053 | ENSG00000017427 | white |
| **COL3A1** | ENSMUSG00000026043 | ENSG00000168542 | white |
| **CCDC80** | ENSMUSG00000022665 | ENSG00000091986 | white |
| **LEP** | ENSMUSG00000059201 | ENSG00000174697 | white |
| **EEPD1** | ENSMUSG00000036611 | ENSG00000122547 | white |
| **NRIP1** | ENSMUSG00000048490 | ENSG00000180530 | white |
| **CCND2** | ENSMUSG00000000184 | ENSG00000118971 | white |
| **EEF2K** | ENSMUSG00000035064 | ENSG00000103319 | white |
| **GADD45A** | ENSMUSG00000036390 | ENSG00000116717 | white |
| **PYGB** | ENSMUSG00000033059 | ENSG00000100994 | white |
| **NDRG1** | ENSMUSG00000005125 | ENSG00000104419 | white |
| **DMRT2** | ENSMUSG00000048138 | ENSG00000173253 | white |
| **LPGAT1** | ENSMUSG00000026623 | ENSG00000123684 | white |
| **QSOX1** | ENSMUSG00000033684 | ENSG00000116260 | white |
| **NUPR1** | ENSMUSG00000030717 | ENSG00000176046 | white |
| **LRP1** | ENSMUSG00000040249 | ENSG00000123384 | white |
| **PIK3R1** | ENSMUSG00000041417 | ENSG00000145675 | white |
| **COL4A2** | ENSMUSG00000031503 | ENSG00000134871 | white |
| **ACVR1C** | ENSMUSG00000026834 | ENSG00000123612 | white |
