## Supplementary material for "Regulatory Modules of Human Thermogenic Adipocytes: Functional Genomics of Meta-Analyses Derived Marker-Genes": Pro FAT genes

Supplementary Table 2. ProFAT Marker-gene list.

| **ProFAT Gene ID** | **Human Code** | **Adipose Tissue marker type** |
| --- | --- | --- |
| UCP1 | ENSG00000109424 | Brown |
| Ndufab1 | ENSG00000004779 | Brown |
| Cpt2 | ENSG00000157184 | Brown |
| Pdha1 | ENSG00000131828 | Brown |
| Shmt1 | ENSG00000176974 | Brown |
| Etfa | ENSG00000140374 | Brown |
| Ndufa8 | ENSG00000119421 | Brown |
| Ndufs3 | ENSG00000213619 | Brown |
| Idh2 | ENSG00000182054 | Brown |
| Pgk1 | ENSG00000102144 | Brown |
| Cs | ENSG00000062485 | Brown |
| Pald1 | ENSG00000107719 | Brown |
| Pgk1-rs7 | ######## | Brown |
| Ndufa9 | ENSG00000139180 | Brown |
| Mecr | ENSG00000116353 | Brown |
| Dnajc15 | ENSG00000120675 | Brown |
| Ppif | ENSG00000108179 | Brown |
| Fahd1 | ENSG00000180185 | Brown |
| Ndufa10 | ENSG00000130414 | Brown |
| Idh3g | ENSG00000067829 | Brown |
| Etfb | ENSG00000105379 | Brown |
| Pkm | ENSG00000067225 | Brown |
| Dld | ENSG00000091140 | Brown |
| Uqcr11 | ENSG00000127540 | Brown |
| Cycs | ENSG00000172115 | Brown |
| Ndufb8 | ENSG00000166136 | Brown |
| Acads | ENSG00000122971 | Brown |
| Gm10053 | ######## | Brown |
| Fh | ENSG00000091483 | Brown |
| Tmem246 | ENSG00000165152 | Brown |
| Sirt3 | ENSG00000142082 | Brown |
| Sdhd | ENSG00000204370 | Brown |
| Slc25a19 | ENSG00000125454 | Brown |
| Ndufs1 | ENSG00000023228 | Brown |
| Cyc1 | ENSG00000179091 | Brown |
| Mrpl34 | ENSG00000130312 | Brown |
| Ndufv1 | ENSG00000167792 | Brown |
| Dlst | ENSG00000119689 | Brown |
| Sdhb | ENSG00000117118 | Brown |
| Etfdh | ENSG00000171503 | Brown |
| Uqcrc1 | ENSG00000010256 | Brown |
| Idh3a | ENSG00000166411 | Brown |
| Mrps36 | ENSG00000134056 | Brown |
| Hadhb | ENSG00000138029 | Brown |
| Gm13910 | ######## | Brown |
| Aco2 | ENSG00000100412 | Brown |
| Slc25a20 | ENSG00000178537 | Brown |
| Impdh1 | ENSG00000106348 | Brown |
| Zic1 | ENSG00000152977 | Brown |
| Acaa2 | ENSG00000167315 | Brown |
| Pdk4 | ENSG00000004799 | Brown |
| Cox7a1 | ENSG00000161281 | Brown |
| Cidea | ENSG00000176194 | Brown |
| Hoxc8 | ENSG00000037965 | White |
| Lpgat1 | ENSG00000123684 | White |
| Gria3 | ENSG00000125675 | White |
| Ar | ENSG00000169083 | White |
| Alcam | ENSG00000170017 | White |
| Sgpp1 | ENSG00000126821 | White |
