## Supplemental Table 3 A, b, C, D for "Regulatory Modules of Human Thermogenic Adipocytes: Functional Genomics of Meta-Analyses Derived Marker-Genes"

**Supplementary Table 3A.**

| **KEGG enriched pathways in BATLAS Brown Marker-genes** | **KEGG enriched pathways in ProFAT Brown Marker-genes** | **KEGG enriched pathways in FTO non-obese Higher expressed genes** |
| --- | --- | --- |
| *Metabolic pathways* | *Metabolic pathways* | *Metabolic pathways* |
| *Thermogenesis* | *Citrate cycle (TCA cycle)* | *Fatty acid metabolism* |
| *Oxidative phosphorylation* | *Carbon metabolism* | *Thermogenesis* |
| *Parkinson's disease* | *Thermogenesis* | *Oxidative phosphorylation* |
| *Huntington's disease* | *Parkinson's disease* | *Carbon metabolism* |
| *Non-alcoholic fatty liver disease (NAFLD)* | *Huntington's disease* | *Huntington's disease* |
| *Alzheimer's disease* | *Oxidative phosphorylation* | *Parkinson's disease* |
| *Carbon metabolism* | *Non-alcoholic fatty liver disease (NAFLD)* | *Fatty acid degradation* |
| *Citrate cycle (TCA cycle)* | *Alzheimer's disease* | *Alzheimer's disease* |
| Retrograde endocannabinoid signaling | Biosynthesis of amino acids | *Non-alcoholic fatty liver disease (NAFLD)* |
| *Fatty acid degradation* | *2-Oxocarboxylic acid metabolism* | Propanoate metabolism |
| *Fatty acid metabolism* | *Fatty acid metabolism* | *Valine, leucine and isoleucine degradation* |
| Propanoate metabolism | Retrograde endocannabinoid signaling | *Citrate cycle (TCA cycle)* |
| *Valine, leucine and isoleucine degradation* | *Pyruvate metabolism* | *PPAR signaling pathway* |
| *Cardiac muscle contraction* | *Fatty acid degradation* | *Fatty acid elongation* |
| Butanoate metabolism | *Valine, leucine and isoleucine degradation* | *Pyruvate metabolism* |
| Lysine degradation | *Fatty acid elongation* | *Cardiac muscle contraction* |
| *PPAR signaling pathway* | *Glyoxylate and dicarboxylate metabolism* | Peroxisome |
| Tryptophan metabolism | Glycolysis / Gluconeogenesis | *Glyoxylate and dicarboxylate metabolism* |
| *2-Oxocarboxylic acid metabolism* | Central carbon metabolism in cancer | Butanoate metabolism |
| *Pyruvate metabolism* | *Cardiac muscle contraction* | *Glycolysis / Gluconeogenesis* |
| Peroxisome | Glycine, serine and threonine metabolism | Adipocytokine signaling pathway |
| *Glyoxylate and dicarboxylate metabolism* | *PPAR signaling pathway* | Tryptophan metabolism |
| Biosynthesis of amino acids |  | Biosynthesis of unsaturated fatty acids |
| beta-Alanine metabolism | | Type I diabetes mellitus |
| Ribosome | | AMPK signaling pathway |
| *Glycolysis / Gluconeogenesis* | | beta-Alanine metabolism |
| Adipocytokine signaling pathway | | Glucagon signaling pathway |
| *Fatty acid elongation* | | Insulin resistance |
|  |  | *2-Oxocarboxylic acid metabolism* |
|  |  | Allograft rejection |
|  |  | Steroid hormone biosynthesis |
|  |  | Graft-versus-host disease |

**Supplementary Table 3B.**

| **REACTOM enriched pathways in BATLAS Brown Marker-genes** | **REACTOM enriched pathways in ProFAT Brown Marker-genes** | **REACTOM enriched pathways in FTO non-obese Higher expressed genes** |
| --- | --- | --- |
| *The citric acid (TCA) cycle and respiratory electron transport* | *The citric acid (TCA) cycle and respiratory electron transport* | *Metabolism* |
| *Metabolism* | *Metabolism* | *Metabolism of lipids* |
| *Respiratory electron transport* | *Pyruvate metabolism and Citric Acid (TCA) cycle* | *The citric acid (TCA) cycle and respiratory electron transport* |
| *Respiratory electron transport, ATP synthesis by chemiosmotic coupling, and heat production by uncoupling proteins.* | *Respiratory electron transport, ATP synthesis by chemiosmotic coupling, and heat production by uncoupling proteins.* | *Fatty acid metabolism* |
| Complex I biogenesis | *Citric acid cycle (TCA cycle)* | *Respiratory electron transport, ATP synthesis by chemiosmotic coupling, and heat production by uncoupling proteins.* |
| *Pyruvate metabolism and Citric Acid (TCA) cycle* | *Respiratory electron transport* | *Respiratory electron transport* |
| *Citric acid cycle (TCA cycle)* | Complex I biogenesis | *Mitochondrial Fatty Acid Beta-Oxidation* |
| Protein localization | *Mitochondrial Fatty Acid Beta-Oxidation* | *Pyruvate metabolism and Citric Acid (TCA) cycle* |
| Mitochondrial translation initiation | *Fatty acid metabolism* | *Signaling by Retinoic Acid* |
| *Fatty acid metabolism* | *Glyoxylate metabolism and glycine degradation* | *mitochondrial fatty acid beta-oxidation of saturated fatty acids* |
| *Mitochondrial Fatty Acid Beta-Oxidation* | *mitochondrial fatty acid beta-oxidation of saturated fatty acids* | *Beta oxidation of hexanoyl-CoA to butanoyl-CoA* |
| *Glyoxylate metabolism and glycine degradation* | *Regulation of pyruvate dehydrogenase (PDH) complex* | *Beta oxidation of decanoyl-CoA to octanoyl-CoA-CoA* |
| Mitochondrial translation elongation | *Beta oxidation of hexanoyl-CoA to butanoyl-CoA* | *Regulation of pyruvate dehydrogenase (PDH) complex* |
| Mitochondrial translation termination | *Beta oxidation of decanoyl-CoA to octanoyl-CoA-CoA* | *Glyoxylate metabolism and glycine degradation* |
| Mitochondrial protein import | *Signaling by Retinoic Acid* | *Citric acid cycle (TCA cycle)* |
| *Metabolism of lipids* | *Metabolism of lipids* | Import of palmitoyl-CoA into the mitochondrial matrix |
| *mitochondrial fatty acid beta-oxidation of saturated fatty acids* | Mitochondrial protein import | Pyruvate metabolism |
| *Beta oxidation of hexanoyl-CoA to butanoyl-CoA* | Lysine catabolism | Beta oxidation of octanoyl-CoA to hexanoyl-CoA |
| Mitochondrial biogenesis | *Import of palmitoyl-CoA into the mitochondrial matrix* | Mitochondrial biogenesis |
| Cristae formation | Metabolism of amino acids and derivatives | Beta oxidation of palmitoyl-CoA to myristoyl-CoA |
| Lysine catabolism | Branched-chain amino acid catabolism | Beta oxidation of lauroyl-CoA to decanoyl-CoA-CoA |
| Beta oxidation of palmitoyl-CoA to myristoyl-CoA | *Transcriptional activation of mitochondrial biogenesis* | Cristae formation |
| Branched-chain amino acid catabolism | Histidine, lysine, phenylalanine, tyrosine, proline and tryptophan catabolism | Peroxisomal protein import |
| Metabolism of amino acids and derivatives |  | Peroxisomal lipid metabolism |
| Beta oxidation of lauroyl-CoA to decanoyl-CoA-CoA |  | Protein localization |
| Beta oxidation of octanoyl-CoA to hexanoyl-CoA | | Endosomal/Vacuolar pathway |
| Beta oxidation of butanoyl-CoA to acetyl-CoA | | Signaling by Nuclear Receptors |
| Pyruvate metabolism |  | Triglyceride metabolism |
| Beta oxidation of decanoyl-CoA to octanoyl-CoA-CoA | | mitochondrial fatty acid beta-oxidation of unsaturated fatty acids |
| Gluconeogenesis |  | Synthesis of very long-chain fatty acyl-CoAs |
| *Signaling by Retinoic Acid* |  | Antigen Presentation: Folding, assembly and peptide loading of class I MHC |
| *Regulation of pyruvate dehydrogenase (PDH) complex* | | Glycerophospholipid biosynthesis |
| Peroxisomal protein import |  | Synthesis of bile acids and bile salts via 24-hydroxycholesterol |
| Peroxisomal lipid metabolism |  | Synthesis of bile acids and bile salts via 27-hydroxycholesterol |
| Mitochondrial calcium ion transport |  | Interferon gamma signaling |
|  |  | Acyl chain remodeling of CL |
|  |  | Formation of ATP by chemiosmotic coupling |
|  |  | Metabolism of steroids |
|  |  | Linoleic acid (LA) metabolism |
|  |  | RA biosynthesis pathway |
|  |  | Synthesis of bile acids and bile salts via 7alpha-hydroxycholesterol |
|  |  | TP53 Regulates Metabolic Genes |
|  |  | Triglyceride catabolism |
|  |  | PPARA activates gene expression |
|  |  | Retinoid metabolism and transport |
|  |  | alpha-linolenic acid (ALA) metabolism |
|  |  | Antigen processing-Cross presentation |
|  |  | Phospholipid metabolism |
|  |  | Visual phototransduction |
|  |  | Triglyceride biosynthesis |
|  |  | Beta oxidation of myristoyl-CoA to lauroyl-CoA |
|  |  | Immunoregulatory interactions between a Lymphoid and a non-Lymphoid cell |

Supplementary Table 3C.

| **KEGG enriched pathways in BATLAS**  **White Marker-genes** | **KEGG enriched pathways in FTO obese Higher expressed genes** |
| --- | --- |
| FoxO signaling pathway | Rap1 signaling pathway |
| AMPK signaling pathway | PI3K-Akt signaling pathway |
| p53 signaling pathway | TGF-beta signaling pathway |
| Focal adhesion | Focal adhesion |
| Glioma | ECM-receptor interaction |
| Melanoma | Malaria |
| AGE-RAGE signaling pathway in diabetic complications | Hippo signaling pathway |
| Amoebiasis | Relaxin signaling pathway |
| Pathways in cancer | Ras signaling pathway |
| Small cell lung cancer |  |
| Relaxin signaling pathway |  |
| PI3K-Akt signaling pathway |  |
| Signaling pathways regulating pluripotency of stem cells | |
| Breast cancer |  |
| Cellular senescence |  |
| Jak-STAT signaling pathway |  |
| Aldosterone-regulated sodium reabsorption |  |
| Transcriptional misregulation in cancer |  |
| Endometrial cancer |  |
| Longevity regulating pathway - multiple species | |
| Non-small cell lung cancer |  |
| Prolactin signaling pathway |  |
| Pancreatic cancer |  |
| Chronic myeloid leukemia |  |
| EGFR tyrosine kinase inhibitor resistance |  |
| Colorectal cancer |  |
| Longevity regulating pathway |  |
| Protein digestion and absorption |  |
| Endocrine resistance |  |
| HIF-1 signaling pathway |  |
| Inflammatory mediator regulation of TRP channels | |
| Progesterone-mediated oocyte maturation |  |
| Human papillomavirus infection |  |
| Prostate cancer |  |
| Insulin resistance |  |
| Cell cycle |  |
| Platelet activation |  |
| Apoptosis |  |
| Insulin signaling pathway |  |
| Measles |  |
| mTOR signaling pathway |  |
| Non-alcoholic fatty liver disease (NAFLD) |  |
| Gastric cancer |  |
| Hepatocellular carcinoma |  |
| Viral carcinogenesis |  |
| Proteoglycans in cancer |  |
| Rap1 signaling pathway |  |

Supplementary Table 3D.

| **REACTOME enriched pathways in BATLAS White Marker-genes** | **REACTOME enriched pathways in FTO obese Higher expressed genes** |
| --- | --- |
| Binding and Uptake of Ligands by Scavenger Receptors | Extracellular matrix organization |
| Signaling by PDGF | Collagen formation |
| Scavenging by Class A Receptors | Diseases of glycosylation |
| Synthesis, secretion, and deacylation of Ghrelin | Elastic fibre formation |
| Collagen degradation | Diseases associated with O-glycosylation of proteins |
| **Signal Transduction** | Collagen biosynthesis and modifying enzymes |
| Assembly of collagen fibrils and other multimeric structures | Integrin cell surface interactions |
| IRS-related events triggered by IGF1R | Assembly of collagen fibrils and other multimeric structures |
| Non-integrin membrane-ECM interactions | Molecules associated with elastic fibres |
| ECM proteoglycans | Defective B3GALTL causes Peters-plus syndrome (PpS) |
| NCAM1 interactions | O-linked glycosylation |
| Signaling by MET | O-glycosylation of TSR domain-containing proteins |
| Platelet activation, signaling and aggregation | Platelet activation, signaling and aggregation |
| Collagen chain trimerization | Degradation of the extracellular matrix |
| Signaling by Receptor Tyrosine Kinases | Collagen chain trimerization |
| **Developmental Biology** | ECM proteoglycans |
| Integrin cell surface interactions | Regulation of IGF transport and uptake by IGFBPs |
|  | Crosslinking of collagen fibrils |
|  | Adherens junctions interactions |
|  | Collagen degradation |
|  | Antagonism of Activin by Follistatin |
|  | Signaling by Activin |
|  | Platelet degranulation |
|  | Post-translational protein phosphorylation |
|  | Syndecan interactions |
|  | Diseases associated with glycosaminoglycan metabolism |
